# MPK12 in stomatal CO_2_ signaling: function beyond its kinase activity

**DOI:** 10.1101/2023.01.31.526436

**Authors:** Chung-Yueh Yeh, Yuh-Shuh Wang, Yohei Takahashi, Katarina Kuusk, Karnelia Paul, Triinu Arjus, Oleksii Yadlos, Julian I. Schroeder, Ivar Ilves, Alfonso T. Garcia-Sosa, Hannes Kollist

## Abstract

- Protein phosphorylation by kinases is a major molecular switch mechanism involved in the regulation of stomatal opening and closure. Previous research defined interaction between MAP kinase 12 and Raf-like kinase HT1 as a required step for stomatal movements by changes in CO_2_ concentration. However, whether MPK12 kinase activity is required for regulation of CO_2_-induced stomatal responses warrants in depth investigation.
- We apply genetic, biochemical, and structural modeling approaches to examining the non-catalytic role of MPK12 in guard cell CO_2_ signaling that relies on allosteric inhibition of HT1.
- We show that CO_2_/HCO_3_^−^-enhanced MPK12 interaction with HT1 is independent of its phosphor-transfer activity. By analyzing gas exchange of plant lines expressing various kinase-dead and constitutively active versions of MPK12 in a plant line where *MPK12* is deleted, we confirmed that CO_2_-dependent stomatal responses rely on MPK12’s ability to bind to HT1 but not its kinase activity. We also demonstrate that purified MPK12 and HT1 proteins form a heterodimer in the presence of CO_2_/HCO_3_^−^ and present structural modeling that explains the MPK12:HT1 interaction interface.
- These data add to the model that MPK12 kinase-activity-independent interaction with HT1 functions as a molecular switch by which stomatal guard cells sense changes in atmospheric CO_2_ concentration.

## Introduction

Carbon dioxide (CO_2_) is not only the main substrate for photosynthesis providing food and fiber to virtually all non-autotrophic organisms, but it is also a major signaling molecule used by many domains of life (Foyer & Noctor, 2020; Hall et al., 2010; Phelan et al., 2021). In plants, CO_2_ controls opening and closure of stomatal pores, which are formed by a pair of guard cells. These highly specialized guard cells can sense both environmental and endogenous signals, and integrate this information to adjust plant gas exchange with the atmosphere. For optimal growth or even survival, plants have to balance the needs of CO_2_ uptake and the risk of losing excess water by transpiration. This is achieved by a guard cell signaling network mainly determined by the crosstalk between CO_2_ and other major factors, such as water availability, light and pathogen attack. Therefore, stomatal responses to changes in CO_2_ concentration are important as they control both water-use efficiency and carbon cycling in ecosystems.

A key event in guard cell CO_2_-sensing involves Mitogen-activated Protein Kinases (MPKs) 12/4 (Marten et al., 2008; Jakobson et al., 2016; Tõldsepp et al., 2018) that can inhibit Raf-like protein kinase High-leaf Temperature 1 (HT1) (Hõrak et al., 2016; Takahashi et al., 2022). HT1 was identified in a screen for mutants that did not respond to changes in CO_2_ concentration (Hashimoto et al., 2006). Plants with recessive or dominant mutations in *HT1* lack stomatal movements in response to CO_2_ changes, but display normal responses to other environmental factors and abscisic acid (Hashimoto-Sugimoto et al., 2016; Hashimoto et al., 2006; Hõrak et al., 2016). Screening natural variants among *Arabidopsis* accessions uncovered MPK12 as an important component of plant water-use and stomatal responses to CO_2_ (Marais et al., 2014; Jakobson et al., 2016). A glycine to arginine substitution (G53R) in the Cvi-0 MPK12 as well as complete deletion of the *MPK12* locus in the Col-0 accession resulted in constitutively more open stomata and reduced stomatal CO_2_ responsiveness (Jakobson et al., 2016). Deletion of *MPK12* combined with guard cell-specific suppression of its close homolog, *MPK4*, abolished CO_2_-dependent stomatal responses, but retained wild-type (wt)-like responses to other environmental factors and abscisic acid (Hõrak et al., 2016; Tõldsepp et al., 2018). Both MPK12 and MPK4 can interact with HT1 and inhibit HT1 kinase activity *in vitro* (Hõrak et al., 2016; Jakobson et al., 2016). As an active kinase, HT1 was capable of phosphorylating the Slow Anion Channel 1 (SLAC1) (Hõrak et al., 2016), Guard cell Hydrogen peroxide-Resistant 1 (GHR1) (Hõrak et al., 2016; Sierla et al., 2018), and two other Raf-like kinases, the Convergence of Blue light and CO_2_ (CBC) 1 and 2 (Hiyama et al., 2017). However, no biological phosphorylated substrate of MPK12 has been found so far, and the purified recombinant MPK12 protein from *E. coli* generally displayed weak autophosphorylation and no detectable transphosphorylation activity (Hõrak et al., 2016; Jakobson et al., 2016; Tõldsepp et al., 2018). MPK12 was shown to phosphorylate artificial substrates only when pulled down from plant cells expressing an epitope-fused MPK12 (Jammes et al., 2009; Lee et al., 2009).

Despite their quintessential roles in phosphor-transfer reactions, MPKs and other protein kinases have recently gained attention in their additional, or sometimes even more prominent, non-catalytic roles in signal transduction (Kung & Jura, 2016; Mace & Murphy, 2021; Matallanas et al., 2011). In most cases, these kinases serve as allosteric regulators to other proteins *via* various mechanisms, whereas some protein kinases were found to modulate transcription by interacting with transcription factors independent of their catalytic activities or even direct binding to DNAs (Hu et al., 2009; Madhani et al., 1997).

A very recent study identified a mechanism of CO_2_ sensing in guard cells in which elevated CO_2_/bicarbonate (HCO_3_^−^) concentration enhanced the interaction of MPK4 and MPK12 with the HT1 kinase, thereby down-regulating HT1-dependent phosphorylation and activation of the downstream kinase CBC1 (Takahashi et al., 2022). A model where MPK12/MPK4:HT1 functions as a CO_2_/HCO_3_^−^ sensor was proposed and supported by mutations in HT1 (Takahashi et al., 2022). Interestingly, a kinase dead MPK12 was shown to be still functional in mediating CO_2_ responses, suggesting a non-catalytic role of MPK12 (Takahashi et al., 2022). However, further research is needed to examine this model of a non-canonical MAP kinase function and to verify MPK12’s mode of action in stomatal CO_2_ signaling.

Here, we apply genetic, biochemical, and structural modeling approaches to investigating the non-catalytic role of MPK12 in the guard cell CO_2_ sensing that relies on allosteric inhibition of HT1. We also demonstrate direct CO_2_/HCO_3_^−^-dependent dimerization of MPK12 and HT1. Furthermore, we show that MPK4, but not MPK11, can complement loss of MPK12 in the guard cells, further supporting the role of MPK12/4:HT1 interaction as a primary CO_2_/HCO_3_^−^ sensor in stomatal guard cells.

## Materials and methods

### Molecular cloning

For protein expression in *Escherichia coli*, coding sequences of *MPK12* (At2g46070.1) variants with a stop codon were PCR-amplified and cloned at the *Bam*HI/*Eco*RI site of pET28-a vector to create 6xHis-tagged recombinant proteins. Point mutations of *MPK12* were introduced by a two-step overlap-PCR. Constructs of His-tagged HT1 (At1g62400.1) and GST-tagged CBC1 (At3g01490.1) were described in Takahashi et al. (2022). For all other constructs used in this study, simplified diagrams can be found in Figure **S1** with more detailed description in the supplementary methods.

### Protein purification from *E. coli* and *in vitro* kinase assay

Recombinant proteins were produced using an *E. coli* protein expression system. Briefly, bacteria were grown in LB or 2xYTQ11 medium at 37°C until the optical density at 600 nm (OD_600_) reached ~0.5 to 0.7, then 0.5 mM isopropyl-β-d-thiogalactopyranoside was added, and *E. coli* were incubated at 20°C for 16 to 24 hours. *E. coli* cells were harvested by centrifugation at 2000g for 20 min and resuspended in Tris-buffered saline [50 mM Tris-HCl (pH 7.5) and 150 mM NaCl]. The *E. coli* cells were disrupted by ultrasonication, and extracted proteins were separated from cell debris by centrifugation at 14,000g for 10 min. GST-tagged proteins and His-tagged proteins were purified using glutathione Sepharose beads and Ni resin beads, respectively.

The *in vitro* kinase assays were performed as follows. For the CBC1 phosphorylation assays, 0.5 μg GST-CBC1, 0.01 μg His-HT1 and 0.5 μg His-MPK12 recombinant proteins were incubated in phosphorylation buffer [50 mM Tris-HCl (pH 7.5), 10 mM MgCl_2_, 0.1% Triton X-100, and 1 mM dithiothreitol (DTT)] with 20 mM NaHCO_3_ or 20 mM NaCl as a control for 30 min, and then proteins were further incubated with 200 μM ATP and 1 μCi [γ-^32^P]-ATP for 30 min. For the MPK12 auto-phosphorylation assays, 2 μg His-MPK12 proteins were incubated in MPK phosphorylation buffer [50 mM tris-HCl (pH 7.5), 10 mM MgCl_2_, 10 mM MnCl_2_, 0.1% Triton X-100, and 1 mM DTT] with 200 μM ATP and 5 μCi [γ-^32^P]-ATP for 60 min.

### Tandem affinity purification of HT1 and MPK12 heterocomplex

The baculovirus containing either His-mCherry-HT1 or twin strep-mVenus-MPK12 was constructed using the Bac-to-Bac™ system (Thermo Scientific). Resulting baculoviruses were used to co-infect 600 ml of Sf9 insect cells in the shaker culture at the density of 2×10^6^ cells/ml at approximate multiplicity of infection 5-10. Cells were grown for 48 hrs, harvested by centrifugation, washed once with ice cold PBS and resuspended in 30 ml of lysis buffer (50 mM Hepes, pH 7.6; 15 mM NaCl; 10% glycerol; 0.5% Tween 20; 10 mM imidazole; 1 mM DTT; 20 mM NaHCO_3_; Roche protease inhibitors cocktail without EDTA). The resulting cell suspension was flash-frozen in liquid nitrogen and stored at −80°C until needed.

To purify the HT1 and MPK12 complex, the cells were thawed and additionally broken in 40 ml Dounce glass homogenizer with 15 strokes of B pestle (all the protein purification steps were carried out either on ice or at 4°C). 5 M NaCl was added to adjust the concentration of this salt to 300 mM and the lysate was cleared by centrifugation at 50000g for 25 min. The cleared lysate was passed through 1 ml of PureCube Ni-NTA agarose column (Cube Biotech). The column was washed three times with N300 (50mM Hepes*NaOH pH 7.6; 300 mM NaCl; 10% glycerol; 0.05% Tween 20; 0.2 mM PMSF; 10 mM imidazole; 1 mM DTT; 20 mM NaHCO_3_) and twice with N100-Ni washing buffer (the same as N300, but with 100 mM NaCl and no PMSF). Proteins were then eluted from the column with N100-Ni buffer + 300 mM imidazole. The eluate was passed through another 0.3 ml Strep-Tactin XT 4Flow column. This was followed by four washes with N100-strep buffer (the same as N100-Ni, but with 25 mM Hepes*NaOH pH 7.6 and no imidazole) and bound protein was eluted from the column with N100-strep + 50 mM biotin buffer. Resulting protein fractions were snap frozen in liquid nitrogen and stored at −80°C.

### Size exclusion chromatography

For the −HCO3- analysis, 400 μg of the tandem affinity purified His-mCherry-HT1 + twin-strep-mVenus-MPK12 protein complex in 140 μl volume was mixed with 400 μl of S200 column buffer (25mM Hepes*NaOH pH 7.6; 100 mM NaCl; 1 mM DTT) and concentrated to approximately 50 μl in Amicon Ultra 0.5 ml Ultracel 10k centrifugation tube. This concentrate was mixed again with 500 μl of S200 buffer and concentrated to 30-40 μl before injecting 30 μl of the final preparation into Superdex 200 5/150 GL size exclusion column. The column was run with S200 buffer at 50 μl/min and 50 μl fractions were collected. Samples from the peak fractions were analyzed by the 10% polyacrylamide–SDS gel electrophoresis and Coomassie brilliant blue staining of the resolved proteins.

For the +HCO3- analysis, the same tandem affinity purified HT1+MPK12 preparation was subjected to the exact same procedure, except that the S200 buffer used in all steps was supplemented with 20 mM NaHCO3.

### Split-ubiquitin yeast-two hybrid (Y2H) assay

The HT1/pDHB1 bait plasmid was transformed into yeast NMY51 (mating type a) while MPK/pPR3-N prey plasmids were transformed into NMY61 (mating type α). For each pairwise interaction assay, pools of prey transformed NMY61 colonies were mated with a single HT1-transformed NMY51 colony, and the mated diploid cells were selected on SD-LeuTrp plates. Ten to 20 colonies were pooled in overnight liquid cultures (SD-LeuTrp) for growth assays. Three μl of serial dilutions (OD_600_=0.5, 0.05, and 0.005) were spotted on SD-LeuTrp and SD-LeuTrpHisAde plates and incubated at 37°C for 3 days and then photographed.

For quantitative β-galactosidase assays, 100 μl of 3 independent pools of diploid yeast cells (OD_600_=1) were mixed with 100 μl of Y-PER reagent:2X β-Galactosidase Assay Buffer (1:1) using the Yeast β-Galactosidase Assay Kit (Thermo Scientific) in a 96-well plate, and OD_414_ were measured with a Multiskan FC device (Thermo Scientific) in a 2-5 min interval for 1-2 hrs. A time point where the highest reading was around 1 was selected for calculating β-galactosidase activities for all samples after subtracting blanks where no yeast cells but culture medium was added in the well. Relative β-galactosidase activities were calculated against the positive control of cells containing HT1/pDHB1 and pAI-Alg5 plasmids.

### Bimolecular fluorescence complementation (BiFC) assays

For ratiometric BiFC, a total of four *Agrobacterium* GV3101 strains each carrying a *X-YFPn*, *Y-YFPc*, *SLAC1-CFP* or *P19* gene silencing suppressor were co-infiltrated to leaves of 4- to 5-week-old *Nicotiana benthamiana* plants at an OD_600_ of 0.02 for each strain in an infiltration buffer (10 mM MgCl_2_, 10 mM MES, pH 5.6, 200 μM acetosyringone). For CO_2_-dependent ratiometric BiFC experiments, infiltrated plants were incubated in an eight-chamber gas-exchange system where each chamber can be set to either high (800 ppm) or low (100 ppm) CO_2_ for 3 days. Fluorescence imaging was acquired with a Zeiss LSM710 confocal microscope equipped with a 20x objective at 3-day post infiltration. Before confocal microscopy, leaves subjected to low CO_2_ treatment were excised and quickly transferred to a portable box where CO_2_ was kept at 100 ppm until imaging. The YFP signals were excited by a 514 nm laser, and emission between 518 and 564 nm was collected. The CFP signals were excited by a 405 nm laser, and emission at 460 to 530 nm was collected. The YFP and CFP signals were collected by sequential scanning to avoid bleed through. Laser power was set to minimize saturation of fluorescence intensity based on the HT1-YFPn and MPK12-YFPc pair, and all parameters were kept identical for the remaining samples.

For the CO_2_-dependent wide field fluorescence microscopy BiFC experiments at UCSD, a single plasmid containing *35S:HT1-NmVen210*, *35S:MPK12-CVen210* and a *P19* silencing suppressor was transformed to *Agrobacterium tumefaciens* GV3101. *N. benthamiana* plants were grown in standard potting soil (Sungrow Horticulture, Professional Growing Mix, MA, USA) under long-day conditions (16-hr-light/8-hr-dark cycle, 22°C) and 60% relative air humidity. Leaves of 5- to 6-week-old plants were infiltrated with agrobacterial inoculants at an OD_600_ of 0.5, which were blinded by a non-coauthor/non-collaborating laboratory member and only unblinded for a third laboratory member after all analyses of blinded data were completed by the experimenter as in Takahashi et al. (2022). For low CO_2_ experiments, plants grown under the above conditions were exposed to low CO_2_ for 2 hrs before infiltration and then incubated in a growth chamber (Percival, IntellusUltra AR-41L2) after infiltration, under low CO_2_ conditions (100 ppm) for 3 days under constant light conditions at 22°C and 60% relative air humidity. For high CO_2_ experiments, the low CO_2_-treated plants were subjected to elevated CO_2_ in a growth chamber for 2 hrs to 800 ppm CO_2_. Microscopy was performed 3 days after infiltration with a Nikon Eclipse E600 fluorescence microscope using a 20x objective lens with an attached INFINITYX digital charge-coupled device color microscopy camera (Takahashi et al., 2022). During microscopy of low CO_2_ samples, plants were gassed continuously with low CO_2_ by passing filtered air through CO_2_ absorbent to maintain the CO_2_ concentration close to nominally 100 ppm. The mVenus signals were excited by a 515 nm laser, and emission between 528 nm was collected. For each leaf, images from non-overlapping areas were captured. For each image, four points from non-overlapping areas were analyzed. Images were obtained and shown using constant imaging conditions. The fluorescence intensity of the images was measured using ImageJ software.

### Western blot analyses

For split-ubiquitin Y2H assays, about 2.5 OD_600_ of yeast cells were harvested by centrifugation and resuspended in 100 μl distilled water and 100 μl of 0.2 M NaOH, incubated for 5 min at room temperature, pelleted, resuspended in 50 μl SDS sample buffer (0.06 M Tris-HCl, pH 6.8, 5% glycerol, 2% SDS, 4% β-mercaptoethanol, 0.0025% bromophenol blue) and boiled for 10 min (Kushnirov, 2000). For ratiometric BiFC experiments, leaf samples (50 mg) were ground under liquid nitrogen and boiled in 100 μl extraction buffer (0.1 M EDTA, pH 8.0, 0.12 M Tris-HCl, pH 6.8, 4% w/v SDS, 10% v/v β-mercaptomethanol, 5% v/v glycerol, 0.005% w/v bromophenol blue) for 10 min (Tsugama et al., 2011). Thirty μl of each sample were separated on a 10% SDS polyacrylamide gel. After electrophoresis, total proteins were transferred onto a nitrocellulose membrane and stained with Ponceau S. Immunodetection of HA-tagged proteins was carried out with a monoclonal anti-HA antibody followed by a goat horseradish peroxidase-conjugated secondary antibody and chemiluminescent detection.

### Plant maintenance and gas-exchange experiments

The *Arabidopsis thaliana MPK12*-deletion (*mpk12-4*) is in the Col-0 accession background, and was isolated previously (Jakobson et al., 2016). Transgenic plants with various binary constructs were generated in the *mpk12-4* background.

For the CO_2_ response phenotyping, seeds were stratified for 2-4 days at 4°C and sown in peat:vermiculite (2:1) mixture in a glass-covered pot to allow separation of rosette leaves from the root and growth media during gas-exchange experiments (Kollist et al., 2007). The plants were cultivated in growth chambers with the following growth conditions: 10/14 hrs light/darkness with 23/19°C and ~60%/80% relative air humidity for light/darkness periods, respectively, and a photosynthetic photon flux density of 250 μmol photons m−2 s−1. Three- to four-week-old plants were analyzed in an eight-chamber gas-exchange measurement system Jyrkki (PlantInvent Ltd.) under conditions similar to the growing period. Plants were allowed to be acclimated in the system under 420 ppm CO_2_ for about 2 hrs until a steady-state stomatal conductance was reached before applying CO_2_ changes. Stomatal conductance was then analyzed in a high (800 ppm)/ambient (420 ppm)/low (100 ppm) CO_2_ regime with duration of 56 min for each treatment.

### Transgenic plant generation and verification

The *Arabidopsis mpk12-4* plants were transformed by the floral dip method (Clough & Bent, 1998). Transgenic seeds were first selected based on the pFAST-R marker (Shimada et al., 2010) that produced red fluorescence in dry seeds. Transgene expression was further verified in the experimental plants’ leaves with a Zeiss LSM710 confocal microscope using a 20x objective. The mVenus signals were excited by a 514 nm laser and the emission was collected in the 518-564 nm wave-length range.

### Statistical Analyses

Statistical analyses were performed using Statistica, version 7.1 (StatSoft) or GraphPad Prism software (version 8.0.1). One-way or two-way ANOVA analyses with Tukey post hoc tests were used, as indicated in figure legends. All effects were considered significant at P< 0.05 unless otherwise stated.

### Structural modeling

Sequences for proteins were obtained from the TAIR website (www.arabidopsis.org). AlphaFold2 deep learning prediction of protein structure was performed with ColabFold using MMseq2, both for monomers and multimers, wild-type and mutants (Mirdita et al., 2021, 2022), without the use of AlphaFold Multimer or templates. Amber-Relax was disabled. A separate multiple sequence alignment (MSA) is generated for each protein. The per-residue confidence metrics (pLDDT) and predicted aligned errors (PAE) parameters of quality of fit were followed and assessed. Models were generated and ranked by per-residue confidence metric. Turbo was used, and a maximum number of sequences was set to 512:1024. Deepmind’s fine-tuned model parameters were used to obtain the PAE per structure, with one trunk of the network run multiple times with different random choices for the MSA cluster centers, three recycles - times the structure is fed back into the neural network for refinement, in addition to one random number seed. Models were superposed for interface comparison.

## Results

### MPK12:HT1 interaction does not require MPK12 kinase activity, but depends on specific amino acids

We previously demonstrated that interaction between MPK12 and HT1 is strongly impaired with a G53R substitution in MPK12, which was identified in the Cvi-0 natural accession of *Arabidopsis thaliana* (Jakobson et al., 2016). *In vitro* kinase assays revealed autophosphorylation activity of MPK12, but not the G53R mutant, implying that loss of kinase activity could affect the interaction. However, a kinase-dead version of MPK12 with arginine substitution to the invariant lysine (K70) required for ATP binding was only slightly impaired in its ability to inhibit HT1 compared to the wtMPK12 and a hyperactive MPK12(Y122C) (Jakobson et al., 2016). These results raised the question of whether MPK12 kinase activity is required for its interaction with HT1. To address this question, we generated another MPK12 mutant with D196G and E200A substitutions, named MPK12(DE). Similar to MPK12(Y122C), MPK12(DE) was designed based on conserved substitutions leading to constitutive active kinases in *Arabidopsis* MPK3/4/6 (Berriri et al., 2012). However, purified recombinant MPK12(DE) showed only weak but comparable autophosphorylation activity to wtMPK12, whereas MPK12(Y122C) yielded a strong autophosphorylation signal in the *in vitro* kinase assay activity (Figure **1a**). On the other hand, the G53R and K70R mutants did not show any detectable autophosphorylation activity (Figure **1a**), consistent with our previous report (Jakobson et al., 2016).

**Figure 1.**
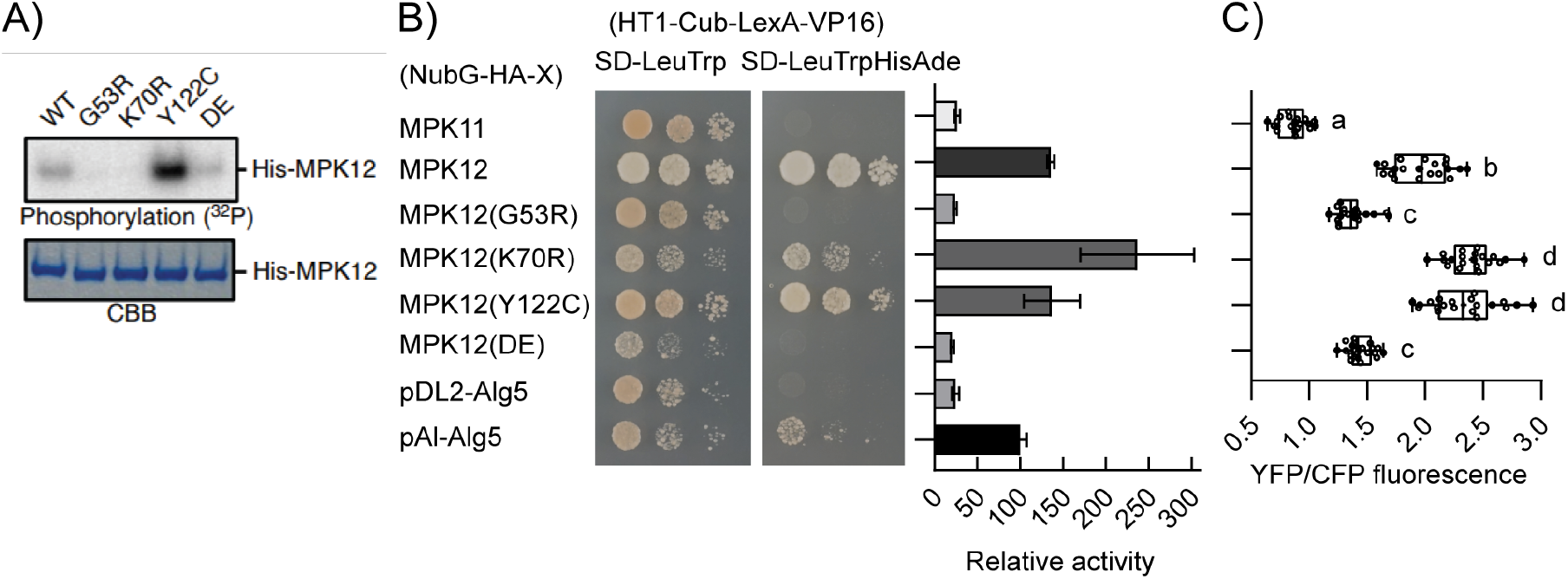
MPK12 interaction with HT1 does not depend on MPK12 kinase activity. **A)** Kinase activity of MPK12 and its variants: WT and DE versions of MPK12 displayed weak autophosphorylation, whereas the Y122C variant had a higher activity; no autophosphorylation activity of the G53R and K70R versions was detected; MPK12 autophosphorylation levels (upper panel) and Coomassie brilliant blue (CBB)-stained gels (lower panel) are shown; the illustrated experiment was repeated 3 times with similar results. **B)** Split-ubiquitin yeast two-hybrid (Y2H) assay: yeast growth with serial dilution on a SD-LeuTrp plate indicates presence of both bait (HT1-Cub-LexA-VP16) and prey (NubG-HA-X) plasmids, and on a SD-LeuTrpHisAde plate to select for interaction; only MPK12, MPK12(K70R) and MPK12(Y122C) interact with HT1; pDL2-Alg5 and pAI-Alg5 are negative and positive prey controls, respectively; bar graph shows relative β-galactosidase activities normalized by the positive control; error bars are SEM (n=3). **C)** Ratiometric bimolecular fluorescence complementation (BiFC) assay in *Nicotiana benthamiana*: box & whisker plot shows stronger interaction between HT1 and MPK12, MPK12(K70R) and MPK12(Y122C); y-axis depicts the split-YFP pairs (HT1-YFPn + MPK-YFPc) used for leaf infiltration, along with *Agrobacterium* carrying a *SLAC1-CFP* construct as a control; x-axis is the ratio of YFP and CFP fluorescence, which indicates interaction strength normalized by the SLAC1-CFP reference control; 20 confocal images taken from three independent leaves were analyzed for each interaction pair; statistically significant groups are denoted with different letters (One-Way ANOVA with Tukey HSD *post hoc* test, P < 0.05).

We next performed interaction assays between HT1 and all five MPK12 variants. We chose the split-ubiquitin yeast-two-hybrid (Y2H) system due to the membrane association nature of HT1 (Hashimoto-Sugimoto et al., 2016; Hõrak et al., 2016). Consistent with our previous report (Jakobson et al., 2016), wtMPK12, but not its close homolog, MPK11, interacted well with HT1. The G53R kinase-deficient MPK12 and the weakly active MPK12(DE) did not interact with HT1, whereas another kinase-dead version, K70R, and the hyperactive Y122C displayed comparable interactions with HT1 as the wtMPK12 (Figures **1b** & **S1a**). To rule out the possibility that lack of interaction was due to poor or no expression of MPK12 proteins in yeast, we performed western analysis and showed that all baits including five MPK12 variants and MPK11 were well expressed (Figure **S2a**).

We also checked the interaction of HT1 and the MPK12 variants in *Nicotiana benthamiana* by the ratiometric bimolecular fluorescence complementation (BiFC) assays. A SLAC1-CFP fusion, which is localized to the plasma membrane where the HT1:MPK12 interaction mainly takes place, was used as a reference control to normalize the BiFC signals (Figure **S1b**). In agreement with the Y2H results, in addition to wtMPK12, the K70R and Y122C variants showed strong interaction with HT1 as indicated by the higher ratio of YFP/CFP fluorescence. On the other hand, the G53R and DE versions of MPK12, as well as MPK11, had very low YFP/CFP signals (Figures **1c** & **S1b**). All split YFP fusion proteins were well expressed as verified by western blot analysis (Figure **S2b**). These data using independent approaches support the model that HT1:MPK12 interaction is independent of MPK12 kinase activity, and the interaction interface may include the G53, D196, and E200 amino acids.

### Kinase activity-independent interaction between MPK12:HT1 depends on CO_2_/HCO_3_^−^ concentration

Both Y2H and BiFC assays allow interaction studies with only a static view due to their irreversibility. Despite the limitation, the irreversible nature of these assays could enhance transient interactions. We modified the BiFC experiments by growing *N. benthamiana* plants under either elevated (800 ppm) or reduced (100 ppm) CO_2_ conditions post-infiltration to capture potential CO_2_-dependent MPK12:HT1 interaction. Using our modified ratiometric BiFC assays, an increase of YFP/CFP signal was observed for wtMPK12 and HT1 under 800 ppm CO_2_ compared to that under 100 ppm. This high CO_2_-induced signal increase was absent for MPK12(G53R), which had weak interaction signals under both high and low CO_2_ conditions (Figure **2a**). Similarly, widefield microscopy-based BiFC assays were conducted with inoculation and subsequent growth at low, 100 ppm, CO_2_, followed by continuous gassing with low CO_2_ air during fluorescence imaging for the low CO_2_-treated samples. These experiments also revealed high CO_2_-induced MPK12:HT1 interaction, which was abolished by the G53R mutation in MPK12 (Figures **2b** & **S1c**). We previously reported that HT1 inhibited SLAC1, whose activation led to stomatal closure, and this inhibition by HT1 was suppressed by MPK12:HT1 interaction (Hõrak et al., 2016). Very recently, we uncovered CO_2_/bicarbonate (HCO_3_^−^)-induced inhibition of CBC1 phosphorylation by HT1 when MPK12 was added to the reaction, and proposed that MPK12 and MPK4, together with HT1 form the primary CO_2_/HCO_3_^−^ sensor in the guard cells (Takahashi et al., 2022). In order to further investigate whether MPK12-induced inhibition of CBC1 phosphorylation by HT1 requires MPK12 kinase activity or only its binding to HT1, we performed *in vitro* kinase assays with the five versions of MPK12 at elevated or low CO_2_/HCO_3_^−^. This experiment showed that 20 mM of CO_2_/HCO_3_^−^ inhibited CBC1 phosphorylation in the presence of HT1 plus wtMPK12, consistent with the recent study (Takahashi et al., 2022). Interestingly, the MPK12(K70R) and MPK12(Y122C) mutant proteins also inhibited CBC1 kinase activity, but not the G53R and DE variants (Figure **2c**). These results support a critical role of MPK12:HT1 interaction in the CO_2_/HCO_3_^−^-sensing mechanism and its independence of MPK12 kinase activity.

**Figure 2.**
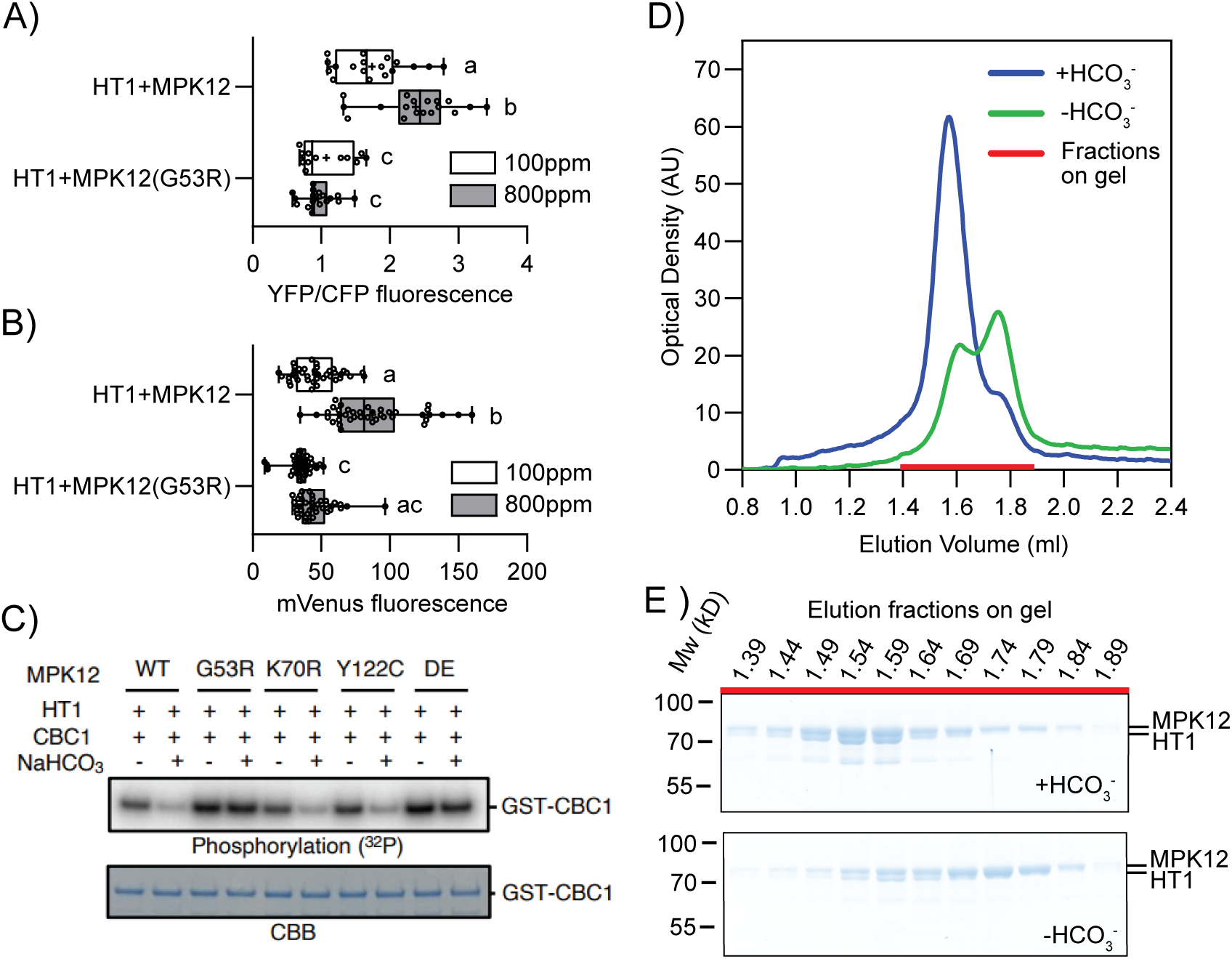
MPK12 interacts with HT1 and inhibits CBC1 phosphorylation by HT1, which are dependent on CO_2_/HCO_3_^−^ concentrations and specific amino acids in MPK12. **A)** Ratiometric confocal microscopy-based BiFC assays in *N. benthamiana* plants kept under reduced (100 ppm) and elevated (800 ppm) CO_2_ conditions after infiltration: MPK12, but not MPK12(G53R), displays increased interaction to HT1 under 800 ppm CO_2_; y-axis indicates split YFP pairs, and x-axis is YFP signal normalized to the SLAC1-CFP reference control; statistically significant groups are denoted with different letters (two-way ANOVA followed by Tukey’s multiple comparisons test; P < 0.005, n=12 for MPK12(G53R) at 100 ppm CO_2_, and 18 for all others). **B)** Widefield microscopy-based BiFC in *N. benthamiana* infiltrated and subsequently grown at 100 ppm CO_2_ for 3 days, and then exposed to 800 ppm or continuing at 100 ppm: MPK12 displays high CO_2_-induced interaction to HT1, which is not evident in the G53R mutant; y-axis shows split mVenus pairs, and x-axis is the mVenus fluorescence intensity; statistically significant groups are denoted with different letters (two-way ANOVA followed by Tukey’s multiple comparisons test; P < 0.05; n=40). **C)** CBC1 phosphorylation by HT1 is reduced by WT, K70R and Y122C versions of MPK12, but not the G53R and DE variants under elevated CO_2_/HCO_3_^−^ condition; His-MPK12s, His-HT1 and GST-CBC1 recombinant proteins were incubated with or without 20 mM NaHCO_3_ for 30 min, and then *in vitro* phosphorylation analyses were performed by adding ^32^P-ATP; CBC1 phosphorylation levels and Coomassie brilliant blue (CBB)-stained gels are shown. **D)** Elution profiles of the Superdex 200 5/150 GL size exclusion chromatography of the His-mCherry-HT1 + twin strep-mVenus-MPK12 protein complex; the same tandem affinity purified protein preparation, purified in the presence of 20mM NaHCO3, was used in both +HCO_3_^−^ and −HCO_3_^−^ runs. **E)**Coomassie brilliant blue stained 10% polyacrylamide-SDS gels showing total proteins in the peak fractions of the chromatography runs depicted in **d)**; the starting elution volumes of the sampled chromatography fractions are indicated on top of each lane; the molecular weight (Mw) standards are indicated on the left; expected Mw of the His-mCherry-HT1 and twin strep-mVenus-MPK12 proteins are 72.9kD and 74.6kD, respectively.

### Purified MPK12 and HT1 form a heterodimer in the presence of bicarbonate

To carry out more direct binding assays for MPK12 interaction with HT1 and its dependence on CO_2_/HCO_3_^−^ concentration, we sought additional systems for protein purification due to difficulties of expressing HT1 in *E. coli*. We co-expressed HT1 and MPK12 proteins in insect cells using the baculovirus expression system followed by tandem purification (Figure **S1d**). During the entire purification process, 20mM of NaHCO_3_ was added to enhance the MPK12:HT1 complex formation.

The purified complex was then subjected to size exclusion chromatography in the presence or absence of added NaHCO_3_. As shown in Figure 2d, in the presence of NaHCO_3_, the main peak is close to the elution volume (Ve) of the 158 kD standard, suggesting that it corresponds to the MPK12:HT1 heterodimer (147.5 KD). This peak is significantly diminished and a second peak with Ve towards the smaller size (75 kD) becomes dominant in conditions without NaHCO_3_, and thus likely corresponds to the mixture of both monomers. The presence of both MPK12 and HT1 in the larger size fraction was confirmed by SDS-PAGE (Figures **2e** & **S3**). These data provide more direct evidence on the CO_2_/HCO_3_^−^-dependent interaction between MPK12 and HT1, and suggest a 1:1 stoichiometry in the MPK12:HT1 complex.

### MPK12 kinase activity is not required for its function in CO_2_-dependent stomatal regulation

Due to the fact that we were unable to detect HT1 phosphorylation by MPK12, it is tempting to speculate that MPK12 exerts its function by sequestering HT1 from phosphorylating downstream effectors independent of the MPK12 phospho-transfer activity. The *mpk12-4* plant, which has the *MPK12* locus completely deleted, together with our five MPK12 variants used in the interaction and kinase assays, provide an excellent platform and tools to address this question. We reasoned that by transforming the *mpk12-4* plants with these MPK12 variants, we could gain insights in the mode of MPK12 function in CO_2_ signaling. We designed and prepared three sets of binary constructs as depicted in Figure S4a. Each set included five constructs of different *MPK12* variants, together with a seed-specific red fluorescent protein marker. The first set contained a 2.5 kb of the *MPK12* genomic sequence without any tag (*gMPK12*) to avoid potential disturbance to the native gene function. The second set had a *mVenus* and a *HA* tag fused to the genomic sequence of *MPK12* (*gMPK12-mVenus-HA*) for easier verification of the guard cell-specific MPK12 expression. The last set aimed to express MPK12-mVenus-HA fusion protein in most cell types by placing the coding sequence between an *Arabidopsis ubiquitin 10* promoter (*pUBQ10*) and a *NOS* terminator (*tNOS*) from the *Agrobacterium tumefaciens* (*pUBQ10:cMPK12-mVenus-HA_tNOS*). All three sets of constructs were used to generate transgenic plants in the *mpk12-4* background. Several T1 seeds with clear red fluorescence were selected from each of the 15 plant lines, and their stomatal responses to CO_2_ changes were examined by a whole plant gas-exchange device (Figure **S4b-d**).

Consistent with previous reports (Jakobson et al., 2016; Tõldsepp et al., 2018), the *mpk12-4* plant had strongly reduced responses to both high and low CO_2_ levels, and its stomatal conductance was about 1.5-fold higher than Col-0 at the ambient CO_2_ (420 ppm) condition (Figure **S4b**). The majority of T1 plants transformed with the wtMPK12 in all 3 sets of constructs displayed Col-0-like stomatal conductance (Figure **S4b**) and responses to both high and low CO_2_ treatments (Figure **S4c,d**), indicating no clear interference to the MPK12 function with the mVenus-HA tag. Among the four MPK12 mutants, only those capable of interacting with HT1 (K70R and Y122C), regardless of their kinase activities, could restore CO_2_ responses in the *mpk12-4* background (Figure **S4c,d**).

We then focused on the set of *gMPK12-mVenus-HA* plants and selected two independent T2 lines from each MPK12 variant for further study. Consistent with the results in T1 plants, only plants with the wtMPK12, K70R and Y122C mutants complemented the *mpk12-4* phenotype with respect to their stomatal CO_2_ responses and the steady state conductance at ambient CO_2_ (Figures **3** & **S5**). Interestingly, despite having impaired CO_2_ responses similar to the *mpk12-4* plants, T2 plants with the G53R and DE versions of MPK12 had lower stomatal conductance than the *mpk12-4* plants (Figures **3a,d** & **S5a,c**). Expression of the MPK12 variants was verified by confocal microscopy based on the mVenus fluorescence, and was found to be guard cell-specific as expected for the native *MPK12* promoter (Figure **3c,f**). All these results clearly suggest a catalytic-independent role of the MPK12 in guard cell CO_2_ signaling.

**Figure 3.**
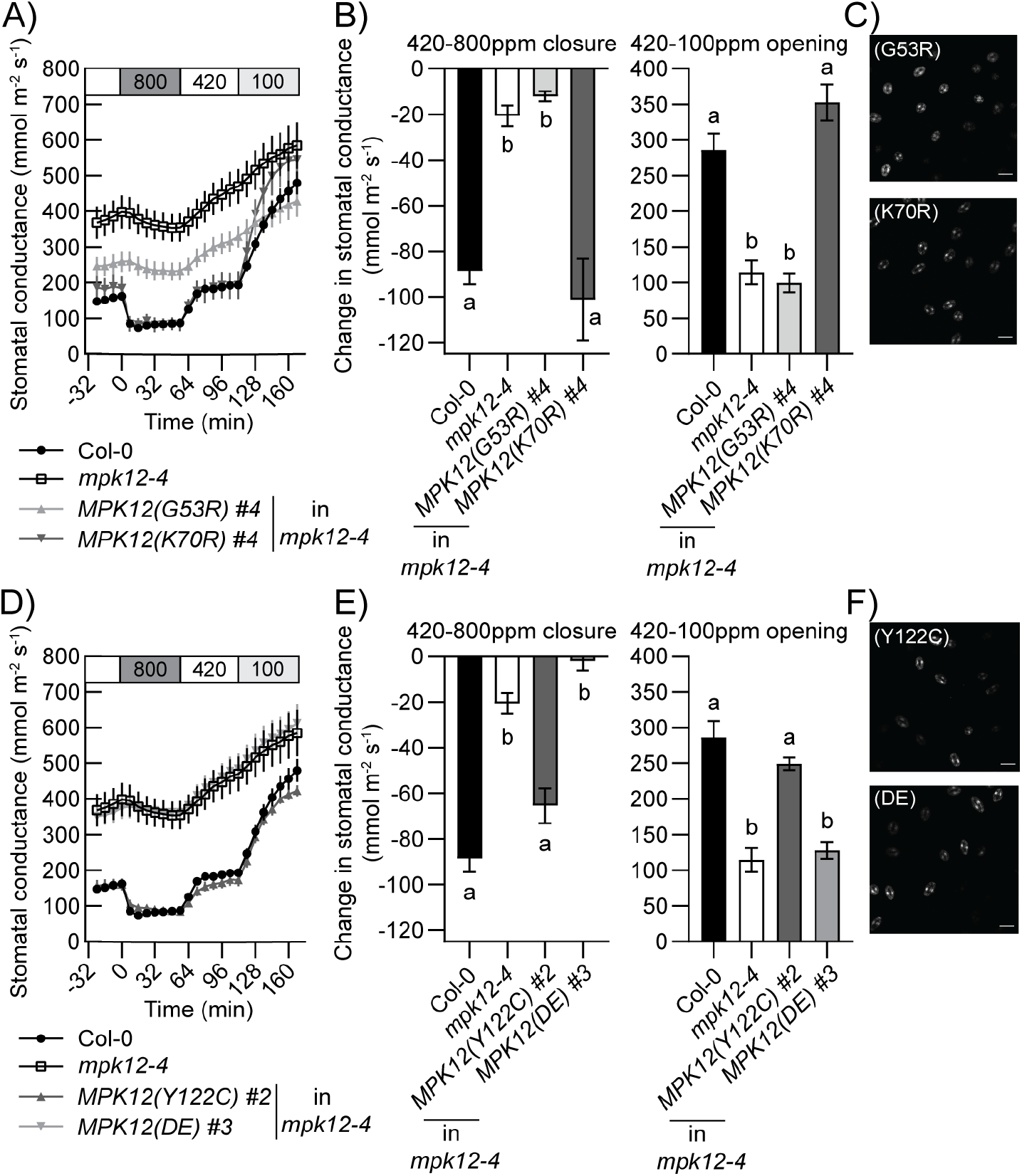
Kinase activity of MPK12 is not required for CO_2_ induced stomatal movements. **A, D)**Time-resolved patterns of whole-plant stomatal conductance in response to changes in CO_2_ concentrations (in parts per million; ppm) as indicated on top of the panels; intact plants of Col-0 (n=13), *MPK12* deletion line *mpk12-4* (n=12), and *mpk12-4* transformed with *MPK12(G53R)* (n=7), *MPK12(K70R)* (n=5) in **a)**and *MPK12(Y122C)* (n=4), *MPK12(DE)* (n=5) in **d)**were analyzed; data of Col-0 and *mpk12-4* controls are the same in **a**& **d** as they were obtained in the same experimental set. **B, E)**Stomatal closure at 16 min after application of 800 ppm CO_2_ (left panels) and stomatal opening after application of 100 ppm CO_2_ for 56 min (right panels); statistically significant groups are denoted with different letters (ANOVA with Tukey HSD unequal N post hoc test, P < 0.05). **C, F)**Representative confocal micrographs of guard cell-specific expression of MPK12 variants fused to mVenus in each transgenic line; scale bar=20 μm.

### A MPK12:HT1 structural modeling reveals important MPK12 amino acids at the interaction interface with HT1

We constructed a predicted complex system of MPK12:HT1 using AlphaFold2 (Jumper et al., 2021). The predicted structure for the MPK12:HT1 complex showed good per-residue confidence metrics, with the residues in the interface area all assessed to have high accuracy (deep blue, Figure **4a**). The predicted aligned errors showed low values for residue pairs of interest, predicting well-defined relative positions. The complex is well-folded, with no overly exposed regions. Important tertiary and secondary structures of MPK12 are recognizable, such as smaller N-lobe and larger C-lobe (Figure **4b**). Residues of MPK12 that are modeled to be important for binding along the interface are G53, D196 (hydrogen bond donated from the side-chain oxygen of D196 to the side-chain nitrogen of K91 in HT1), and E200 (hydrogen bond from the backbone oxygen of E200 to the side-chain NH_2_ of R102 in HT1), along with residues in the substrate binding region: N263, Y277, and P285. Deletion of D196, E200, and G53 resulted in a predicted unfavorable binding due to loss of intermolecular contacts, as well as to MPK12 function. On the other hand, residues K70 and Y122 are predicted not to be in the contact surface between MPK12 and HT1 (Figure **4c**), and their mutation also does not affect physiological functions (Figures **3**, **S4** & **S5**).

**Figure 4.**
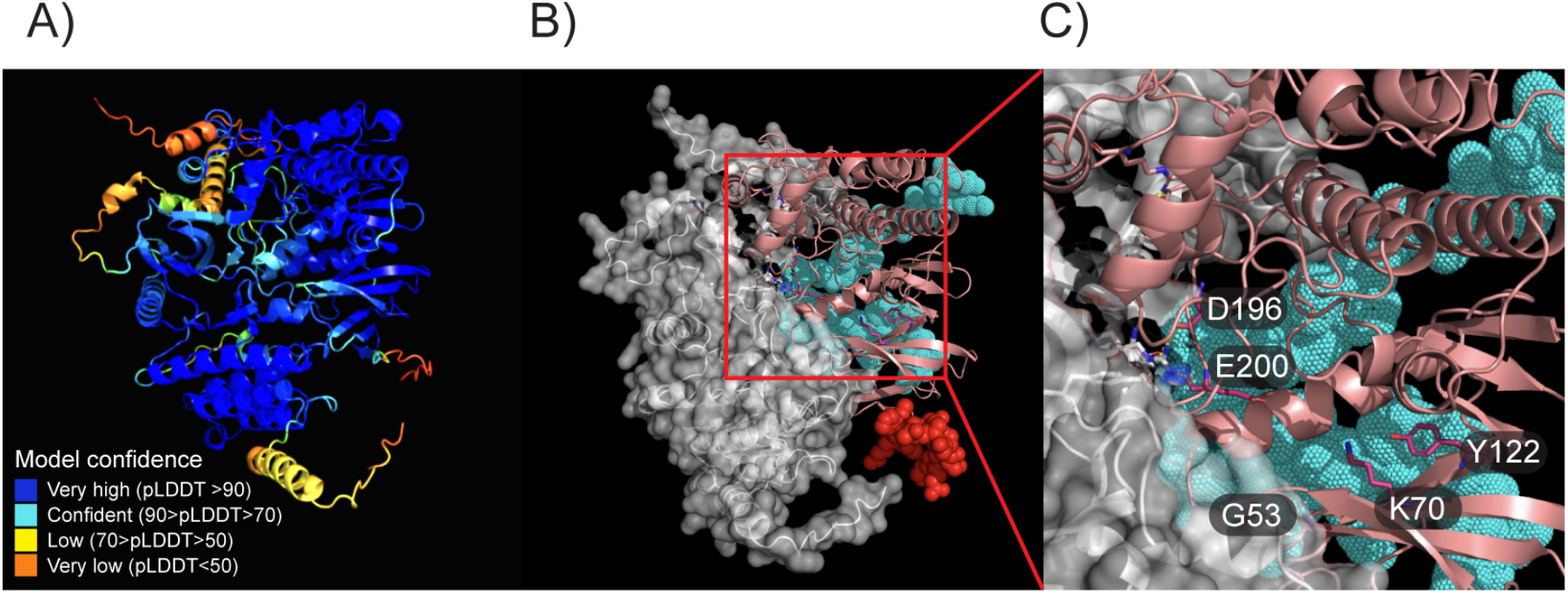
AlphaFold2 structural prediction model for the complex of MPK12 with HT1. **A)** Color-coded model confidence indicates a very good overall confidence, with very high accuracy at the interface area indicated by the per-residue confidence metrics (pLDDT). **B)** HT1 (gray, surface-filled structure) in complex with MPK12 (pink, ribbon structure) with highlights in the N-terminal region (15 amino acids, red spheres), C-terminal region (41 amino acids, cyan dots), and residues G53, K70, Y122, D196, and E200 of MPK12. **C)** Closeup of the interaction interface highlighting MPK12 interface residues G53, D196, and E200, and MPK12 non-interface residues K70 and Y122, of MPK12.

### MPK4, but not MPK11, complements *MPK12*-deletion phenotype

There are 20 *MPK*s in the *Arabidopsis* genome, which can be further separated into four (A to D) subgroups based on sequence homology. *MPK12* is most highly similar to *MPK4* and *MPK11* in the subgroup B (Hamel et al., 2006; Ichimura et al., 2002). We have previously shown that *MPK12* is diverged from *MPK4*, and is specific to the Brassicaceae among angiosperms (Tõldsepp et al., 2018). *MPK12* is principally expressed in the guard cells, whereas *MPK4* is more ubiquitous and has broader functions. Accordingly, deletion of the *MPK12* gene does not affect plant growth, whereas the *MPK4* knockout mutants are extremely dwarfed (Petersen et al., 2000; Tõldsepp et al., 2018). Expressing *MPK12* under the *MPK4* promoter could not rescue the dwarfism phenotype, indicating MPK12 has evolved as a more specialized stomatal regulator at the protein level (Tõldsepp et al., 2018). However, it has not been tested whether the more ancient MPK4 or other close homologs, such as MPK11, can substitute the MPK12 function in the guard cells. We fused the coding sequence of *MPK4* or *MPK11* to *mVenus-HA*, and placed it under the control of *MPK12* promoter and terminator. All plants transformed with the *MPK4* construct which displayed guard cell-specific mVenus fluorescence restored the CO_2_ responses in the *mpk12-4* background. On the contrary, none of the *MPK11* transgenic plants, despite having clear guard cell mVenus fluorescence, could complement the *mpk12-4* phenotype (Figures **5** & **S6**).

**Figure 5.**
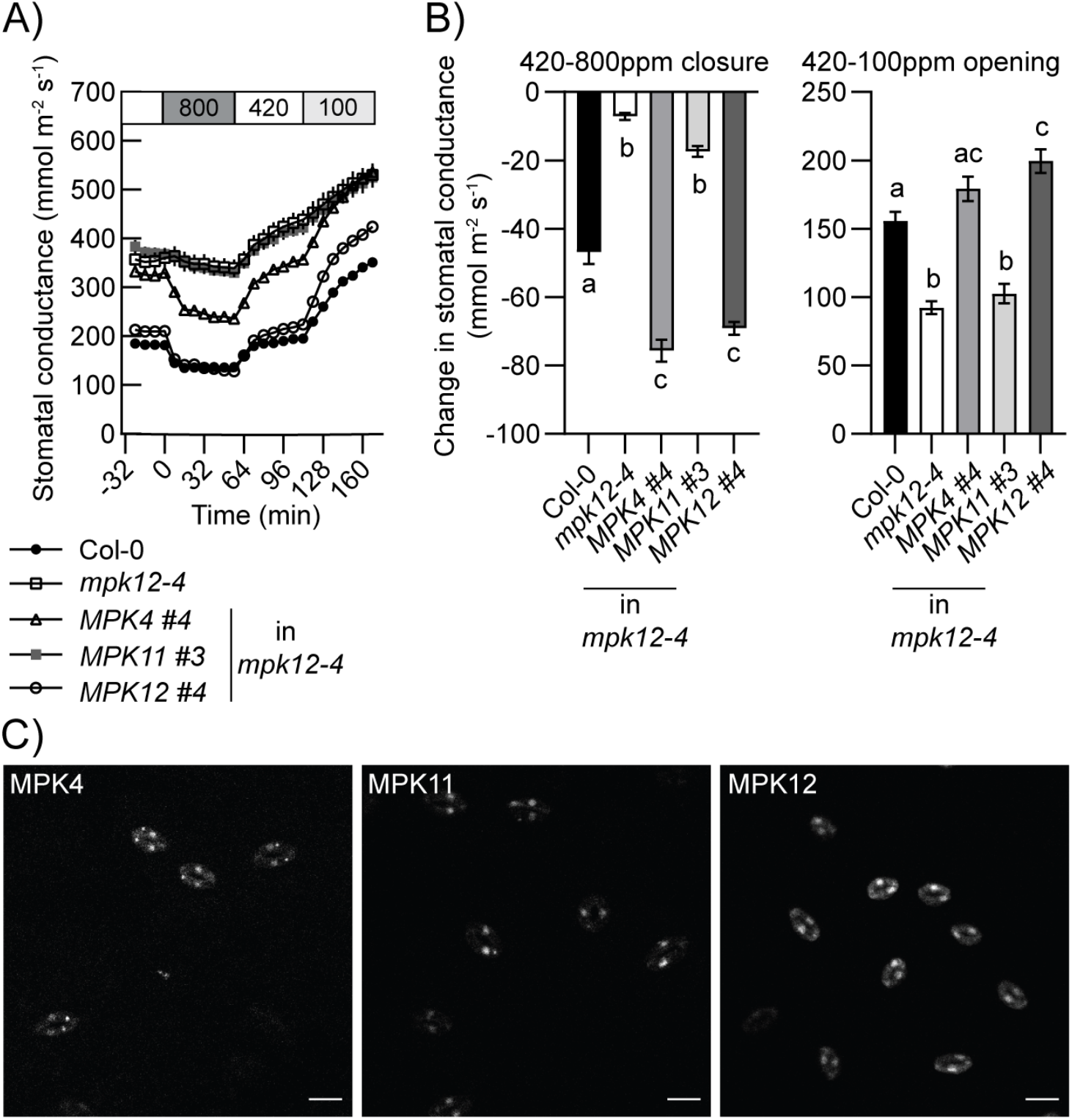
Expression of MPK4 in guard cells can restore *mpk12-4* CO_2_ responses. **A)** Time-resolved patterns of stomatal conductance in response to changes in CO_2_ concentrations (in parts per million; ppm) as indicated on top of the panels of *mpk12-4* plants complemented with *pMPK12:MPK4/11/12-mVenus-HA* constructs and Col-0 plants (n=4-5). **B)** Stomatal closure at 16 min after application of 800 ppm CO_2_ (left panels) and stomatal opening after application of 100 ppm CO_2_ for 56 min (right panels); statistically significant groups are denoted with different letters (ANOVA with Tukey HSD unequal N post hoc test, P < 0.05). **C)** Representative confocal micrographs of guard cell-specific expression of MPK-mVenus fusion; scale bar=20 μm.

## Discussion

In the present study, we characterize a function of MPK12 in stomatal CO_2_ sensing that is independent of its kinase activity, but requires interaction with the HT1 in a CO_2_/HCO_3_^−^-dependent manner. Our results with point mutations in MPK12 (Figures **1-3** & **S5**), along with recent findings by Takahashi et al. (2022), strongly suggest MPK12 as an allosteric inhibitor for the HT1 kinase. Together these two proteins form a dynamic complex that responds to the intercellular CO_2_/HCO_3_^−^ concentration (Figure **2**; Takahashi et al., 2022) and thus modulate stomatal CO_2_ responses.

In support of our model that MPK12:HT1 regulates stomatal movements *via* complex formation or dissociation depending on the CO_2_ levels, we demonstrate that the phospho-transferase activity of MPK12 can be uncoupled from its ability to interact with HT1 (Figure **1**). The MPK12(G53R) variant was identified from the natural Cvi-0 accession of *Arabidopsis thaliana*, and found to be defective in its kinase activity as well as ability to interact with HT1. The G53 residue is located in the glycine-rich loop (GxGxx**G**) between β1 and β2 in the N-lobe of the bi-lobed protein kinase architecture (Figure **4c**). This loop is the most flexible part in the N-lobe, and is responsible for positioning the γ-phosphate of ATP for catalysis. Another variant, MPK12(K70R), was created by mutating the conserved lysine in the Ax**K** motif in β3, which is also part of the ATP-binding pocket. Substitutions in these regions are frequently found in pseudokinases, and mutations in active kinases normally resulted in kinase deficiency (Taylor & Kornev, 2011). We confirm loss of autophosphorylation in the G53R and K70R versions of MPK12 (Figure **1a**; Jakobson et al., 2016). On the other hand, we generated two versions of “hyperactive” MPK12 based on mutations in corresponding residues that resulted in elevated basal kinase activities of the *Arabidopsis* MPK3/4/6 in yeasts (Berriri et al., 2012). The MPK12(Y122C) has a tyrosine to cysteine conversion at the gatekeeper residue that connects two hydrophobic spines, the regulatory (R) and catalytic (C) spines, and thus regulates ATP binding. Replacing this residue with a bulkier amino acid tends to stabilize the R-spine and favors kinase activation (Taylor & Kornev, 2011). However, in one of the mammalian MPKs, the ERK2, mutations of the gatekeeper glutamine (Q103) to a smaller glycine or alanine increased basal activity due to autoactivation, which was attributed to higher backbone flexibility in the activation lip (Emirick et al., 2006). The conversion from a bulkier tyrosine to smaller cysteine in the Y122C mutant of MPK12 may also lead to autoactivation via a similar mechanism. The DE mutants contained substitutions in two amino acids, and this version of *Arabidopsis* MPK4 exhibited MAP2K-independent activity *in planta*, whereas MPK4(Y124C) was hyperactive when purified from *E. coli*, but not *in planta* (Berriri et al., 2012). Our kinase assay results revealed strong autophosphorylation in the recombinant MPK12(Y122C) purified from *E. coli*, but only weak phosphorylation in MPK12(DE) (Figure **1a**). However, whether MPK12(DE) is hyperactive *in planta* requires further study by purifying proteins from the transgenic plants.

In total, we generated five amino acid mutations which are distributed across various conserved domains of the MPK12 and have differential effects on the kinase activity. Our interaction study and *in vitro* kinase assays clearly demonstrate that interaction between MPK12 and HT1 can be induced or enhanced by elevated CO_2_ or HCO_3_^−^, and does not require kinase activity of MPK12 (Figures **1** & **2**). Indeed, a computer-based structural modeling maps G53, D196 and E200 in MPK12 to the interface of the MPK12:HT1 complex, whereas MPK12 residues of K70 and Y122 are predicted to be away from the interface (Figure **4**). Structural modeling has also suggested that strong dominant mutations in the HT1 protein lie at the predicted interface of MPK4/MPK12 with HT1 (Takahashi et al., 2022). Together these models, supported by CO_2_-induced stomatal movement phenotypes of respective mutant plants, implicate the relevance of the MPK12 and HT1 interface for functional CO_2_ signaling. High CO_2_ concentration may potentially produce changes in binding due to pH modification. These could be changes in ionization states of polar amino acids, as well as chemical modification of specific side chains that could produce inter-residue links.

We reasoned that these MPK12 variants, each displaying contrasting features with respect to kinase activity and ability to interact with HT1, could be useful tools in addressing the question: Is MPK12-mediated stomatal CO_2_ signaling dependent on the MPK12 catalytic activity or its ability to modulate HT1 *via* allosteric inhibition, or even both? Using the *MPK12*-deletion (*mpk12-4*) line as a platform, we demonstrate full restoration of CO_2_ responses in the *mpk12-4* plants complemented with the wtMPK12, MPK12(Y122C), and more importantly, the kinase-dead MPK12(K70R). On the other hand, MPK12(G53R) and MPK12(DE) could not rescue *mpk12-4* CO_2_-insensitive phenotype (Figures **3**, **S4** & **S5**). These results are in accordance with the interaction study (Figure **1**) and kinase assays (Figure **2c**), supporting a non-catalytic role of MPK12 in guard cell CO_2_ signaling.

Eukaryotic protein kinases are enzymes with a highly conserved bilobed protein fold capable of phosphor-transfer *in trans* or *in cis* (*i.e*. transphosphorylation or autophosphorylation). However, unlike other catalytic enzymes, they are very inefficient in terms of catalysis. All protein kinases transition between inactive and active states, yet eukaryotic protein kinases seem to shift the equilibrium mostly towards the inactive state, and only become active transiently under specific conditions (Taylor & Kornev, 2011). Eukaryotic protein kinases often function as molecular switches, resembling the GTP-binding proteins, which regulate a diverse array of biological processes by switching between “on” and “off” states upon GTP binding (Shaw et al., 2014). By acting sequentially, such as in the case of the MAPK cascades which comprise of at least three tiers of kinases, even a weak initial signal can be amplified and transduced to the targeted site. However, compelling evidence has emerged that kinases can carry out important functions beyond phosphorylation. A growing number of mammalian and yeast kinases have been demonstrated to play non-catalytic roles such as being allosteric regulators or scaffolds for specific protein assemblies. The MAP kinase ERK2 was shown to regulate transcription by directly binding to DNA through a cluster of positively-charged amino acids in the C lobe of the kinase domain (Hu et al., 2009). However, no such non-catalytic functions of active plant kinases, except for some instances in pseudokinases (Sierla et al., 2018), has been reported *in vivo*. Our findings, together with Takahashi et al. (2022), represent a founder example of a plant MAP kinase that plays an important function independent of its catalytic activity.

We further show that two close homologs of MPK12 display distinct abilities to interact with HT1, and this ability is in accordance to whether they are capable of complementing loss of *MPK12 in planta*. Among the five *Arabidopsis* group B MPKs, MPK4, 11 and 12 share highest homology, yet only MPK4 and 12 are able to inhibit HT1 *via* protein-protein interaction (Hõrak et al., 2016; Jakobson et al., 2016; Takahashi et al., 2022). Accordingly, *MPK4*, but not *MPK11*, when driven under the *MPK12* promoter, can restore CO_2_ responses in the *mpk12-4* plant (Figures **5** & **S6**). These results are consistent with findings that MPK11 did not enable high CO_2_-triggered CBC1 kinase activation *in vitro* and that MPK11 did not interact with HT1 (Takahashi et al., 2022; Jakobson et al., 2016). Furthermore, these data support a hypothesis that MPK12 is diverged from MPK4, which possesses broader functions and a more ubiquitous expression pattern. Therefore, MPK4 can substitute MPK12, but not *vice versa*, when expressed under specific promoters (Figure **5**).

## Acknowledgements

This research was supported by Estonian Research Council (Grants PRG430 to H.K., PRG1509 to A.T.G.-S. and PRG757 to I.I.), European Regional Development Fund (Centre of Excellence in Molecular Cell Engineering (CEMCE) to H.K. & Y.-S.W., JST, PRESTO Grant Number JPMJPR21D8 to Y.T. and a National Science Foundation grant (MCB-1900567) to J.I.S.

## Author contributions

Y.-S.W. and H.K. conceived the study. Y.-S.W., A.T.G.-S and I.I. designed the research. Y.-S. W., C.-Y. Y., Y.T., K.K., K.P., T.A., O.Y., I.I., and A.T.G.-S. conducted the experiments. Y.-S. W., C.-Y. Y., Y.T., K.K., K.P., T.A., I.I., J.I.S, A.T.G.-S. and H.K. analyzed the data. Y.-S.W. and H.K. wrote the manuscript. All authors read and approved the manuscript.

## Supporting information

### Supplementary methods: Molecular cloning

For split-ubiquitin yeast two-hybrid (Y2H) assays, the DUALhunter kit (Dualsystems Biotech) was used for bait and prey construction. The *HT1* coding sequence without a stop codon was cloned between two *Sfi*I sites in the pDHB1 bait vector to allow in-frame fusion with the *Cub-LexA-VP16*. For prey constructs, coding sequences of *MPK12* variants and *MPK11* (At1g01560.2) were fused to *NubG-HA* in the pPR3-N vector. Plasmids of pAI-Alg5 (with a native NubI) and pDL2-Alg5 (with a mutated NubG having I13G substitution to prevent spontaneous reconstitution of split ubiquitin) were used as positive and negative prey controls, respectively (Figure **S1a**).

For bimolecular fluorescence complementation (BiFC) assays, two different sets of binary constructs were used. In the ratiometric confocal microscopy-based BiFC experiments, all binary constructs were created by introducing gene of interest (*HT1* and *MPKs*) via ligation-independent cloning into a *35S:YFPn* (amino acids 1-173 of eYFP) or *35S:YFPc* (amino acids 155-239 of eYFP) vector as describe in Hõrak et al., 2016. A *35S:SLAC1-CFP* construct was used as a transformation control and a ratiometric reference marker (Figure **S1b**; Hõrak et al., 2016). For the widefield CO_2_-dependent BiFC experiments, a modified split mVenus system with a splitting point at the 210th residue (Gookin & Assmann, 2014) was used to reduce background signals. Binary constructs with dual expression cassettes of the *35S:HT1-NmVen210* (amino acids 1-210 of mVenus) and *35S:MPK-CVen210* (amino acids 211-239 of mVenus) were sequentially assembled by Golden Gate cloning. First, the *35S:HT1-NmVen210* cassette was assembled into a destination vector, P19_amilCP/pYSWb1, which contained a *P19* silencing suppressor and an *amilCP* dropout. The *amilCP* dropout was flanked by two outward-oriented *Bsa*I sites to allow replacement of the purple chromoprotein with the desired expression cassette by Golden Gate cloning with a GGAG-CGCT linker according to the common syntax for plants (Patron et al., 2015). After introduction of the *35S:HT1-NmVen210* cassette, another *amilCP* dropout digested with *Sgs*I and *Sal*I was added at the downstream *Mlu*I/*Sal*I site to allow assembly of the *35S:MPK-CVen210* cassette (Figure **S1c**).

For *in planta* complementation, four sets of *MPK* cassettes were assembled by Golden Gate cloning in the destination vector, pFAST-R_amilCP/pYSWb1, which contains a *pFAST-R* marker (Shimada et al., 2010) and an *amilCP* dropout for Golden Gate assembly, as well as a *P19* silencing suppressor and a hygromycin-resistance cassette (*HygR*). The first set (*pMPK12:gMPK12_tMPK12*) contained a 2.7 kb genomic sequence of *MPK12* encompassing 543 bp upstream of the start codon (*pMPK12*) and 520 bp downstream of the stop codon (*tMPK12*). The second set (*pMPK12:gMPK12-mVenus-HA_tMPK12*) contained the 2.7 kb genomic *MPK12* fused to a *mVenus-HA* tag. The third set (*pUBQ10:cMPK12-mVenus-HA_tNOS*) had a cDNA copy of *MPK12* fused to *mVenus-HA* driven by an *Arabidopsis ubiquitin10* promoter (*pUBQ10*) and the *nopaline synthase* terminator (*tNOS*) from *Agrobacterium tumefaciens*. Each of the three sets include MPK12 from the wild-type Columbia-0 (Col-0) and four variants with 1 or 2 amino acid substitutions, which were created by overlap-PCR. The fourth set (*pMPK12:cMPK-mVenus-HA_tMPK12*) had a cDNA copy of various MPKs (MPK4, 11, 12 and chimeras/truncations) fused to a *mVenus-HA* tag under the control of *MPK12* regulatory elements (Figure **S4a**).

For protein expression in the baculovirus/insect cell system, the HT1 cDNA was fused with a N-terminal His tag (MHHHHHH; 6xHis following a methionine), mCherry and a TEV protease cleavage site. The entire coding sequence was cloned at the *Sal*I/*Bcu*I site of the pFastBac1 vector (Thermo Scientific). Similarly, the MPK12 cDNA was N-terminally fused with a twin strep tag followed by a HA epitope and mVenus (Figure **S1d**).

**Figure S1.**
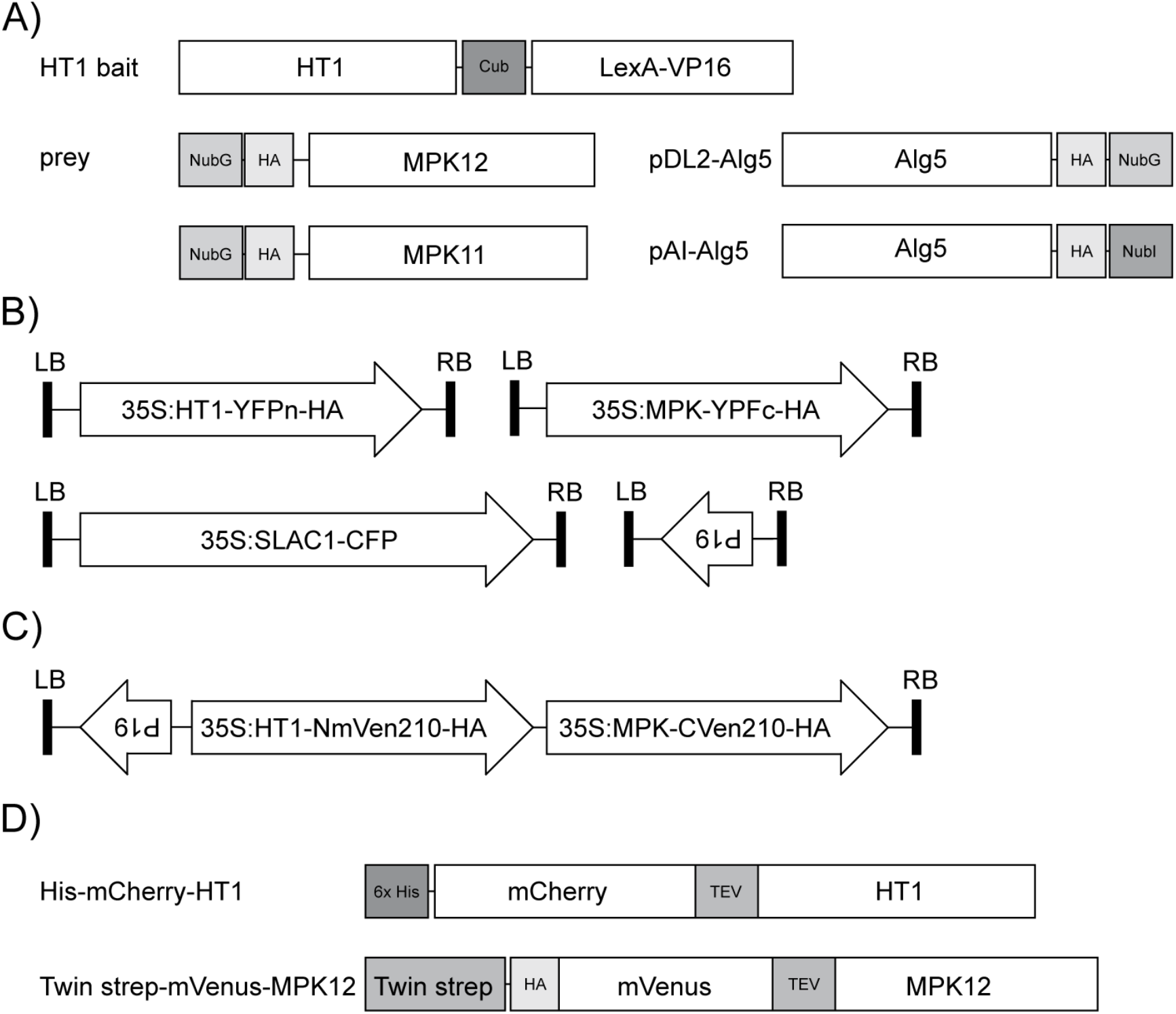
Construct schemes used in this study. **A)** Diagram of constructs used in split-ubiquitin Y2H assays: HT1, as a bait, is fused to the C-terminal half of ubiquitin (Cub) and a transcription factor (LexA-VP16); MPK12s and MPK11 are fused to a mutated N-terminal half of ubiquitin with an I13G substitution (NubG) and a hemagglutinin (HA) tag; a yeast Alg5 coding sequence fused to an HA tag and NubG (pDL2-Alg5) is used as a negative prey control, whereas Alg5 fused to HA and the native N-terminal ubiquitin (NubI) is used as a positive control (pAI-Alg5). **B)** Diagram of constructs used in ratiometric BiFC assay: the T-DNA region of four separate binary constructs are shown; each contains either a P19 silencing suppressor, a HT1 fused to the N-terminal half (1-173 amino acids) of eYFP and a HA tag (HT1-YFPn-HA), a MPK fused to the C-terminal half (155-239 amino acids) of eYFP and a HA tag (MPK-YFPc-HA), or a plasma membrane-localized Slow Anion Channel 1 (SLAC1) fused to eCFP as a transformation and reference control. **C)** Diagram of constructs used in the widefield fluorescence microscopy-based BiFC assay: the T-DNA region contains a P19 suppressor and two split mVenus fusions (split between 210th and 211th amino acids) with a HA tag (HT1-NmVen210-HA & MPK-CVen210-HA) for interaction study; all three cassettes are in one plasmid to ensure coexpression. **D)** Diagram of constructs used for recombinant protein expression using the baculovirus/insect cell system. The HT1 construct contains a 6xHis tag immediately following the methionine, as well as a mCherry fluorescent protein and a TEV protease cleavage site, whereas the MPK12 construct has a twin strep tag followed by a HA epitope, mVenus and a TEV site.

**Figure S2.**
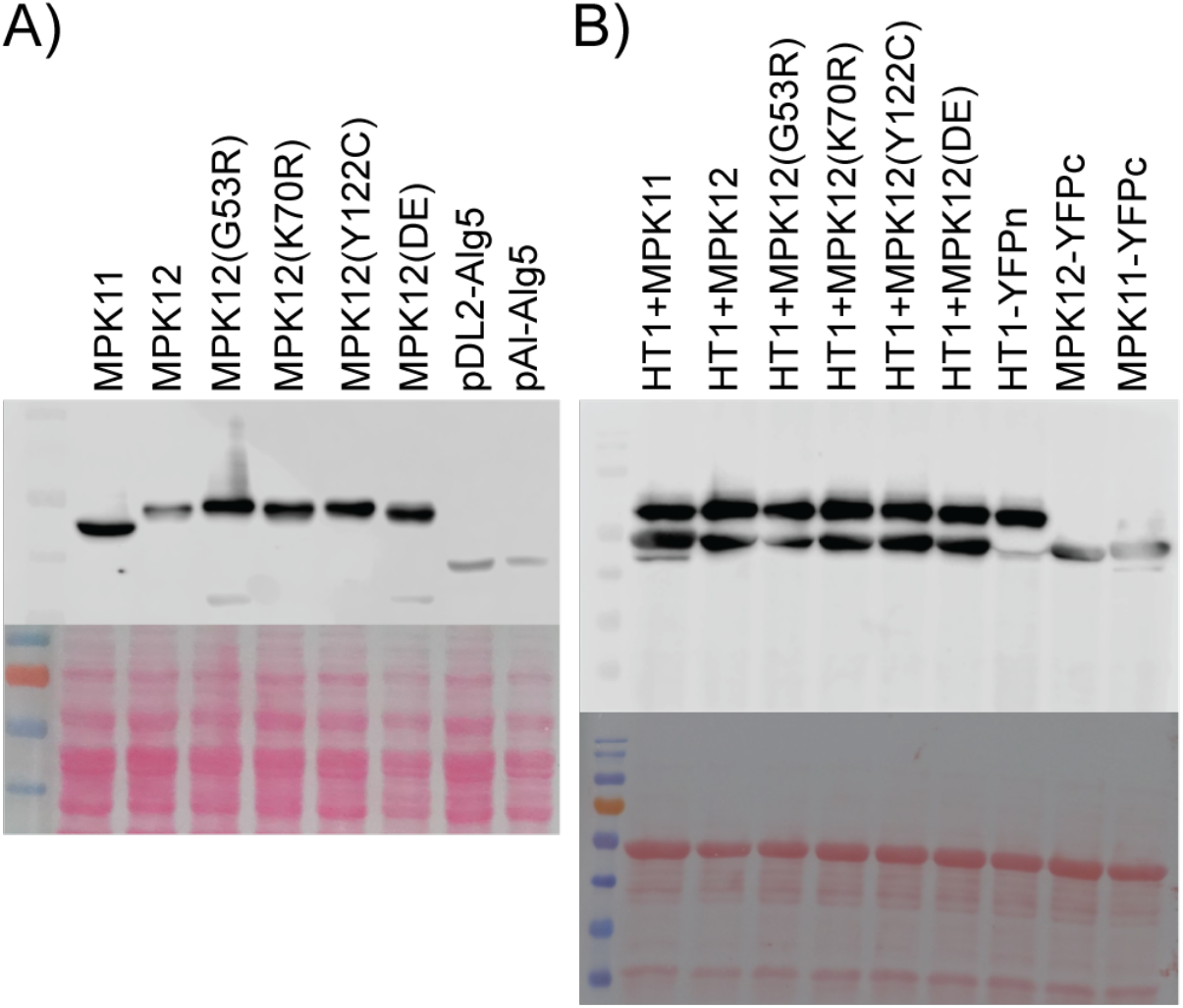
Western blot analyses indicate successful expression of all fusion proteins in the interaction studies. **A)** Western blot analyses with anti-HA antibody (upper panels) and Ponceau S staining (lower panels) of proteins extracted from yeast cells in the Y2H study show expression of all HA-fusion proteins from the prey constructs; no signals from HT1 due to lack of a HA tag in the bait construct. **B)** Western blot analyses with anti-HA antibody (upper panels) and Ponceau S staining (lower panels) of proteins extracted from *Nicotiana benthamiana* leaves in the ratiometric BiFC experiments show expression of HT1-YFPn and MPK-YFPc proteins; proteins extracted from leaves infiltrated with *SLAC1-CFP* plus *HT1-YFPn*, *MPK12-YFPc* or *MPK11-YFPc* are used as size controls for HT1 and MPK fusions.

**Figure S3.**
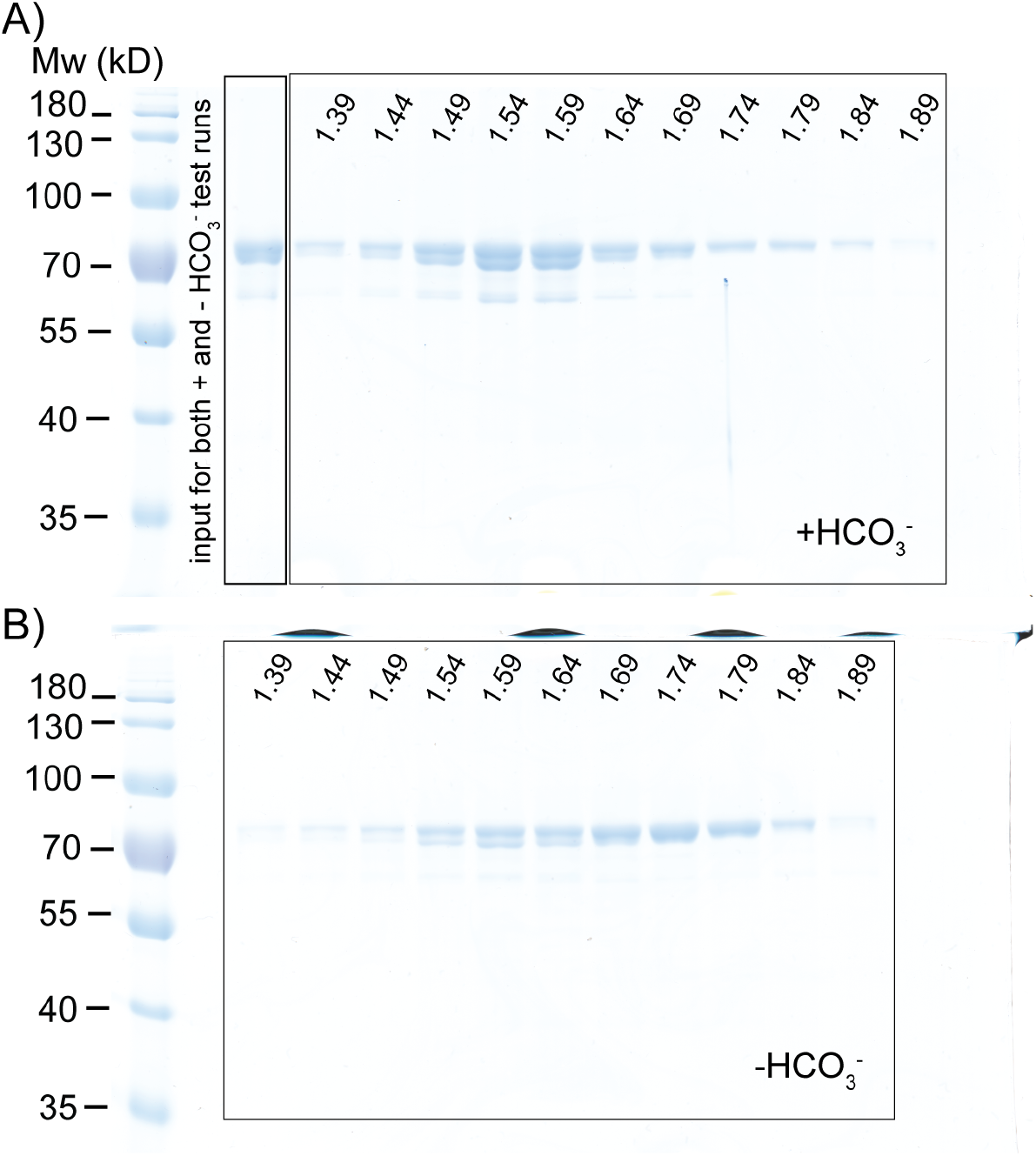
Whole gels of protein fractions shown in Fig. 2e. Coomassie brilliant blue stained 10% polyacrylamide-SDS gels showing total proteins in the peak fractions of the chromatography runs with HCO_3_^−^ in **a)**and without added HCO_3_^−^ in **b)**as depicted in **Fig. 2e**; the starting elution volumes of the sampled chromatography fractions are indicated on top of each lane; the molecular weight (Mw) standards in these gels are indicated on the left; expected Mw of the His-mCherry-HT1 and twin strep-mVenus-MPK12 proteins are 72.9kD and 74.6kD, respectively.

**Figure S4.**
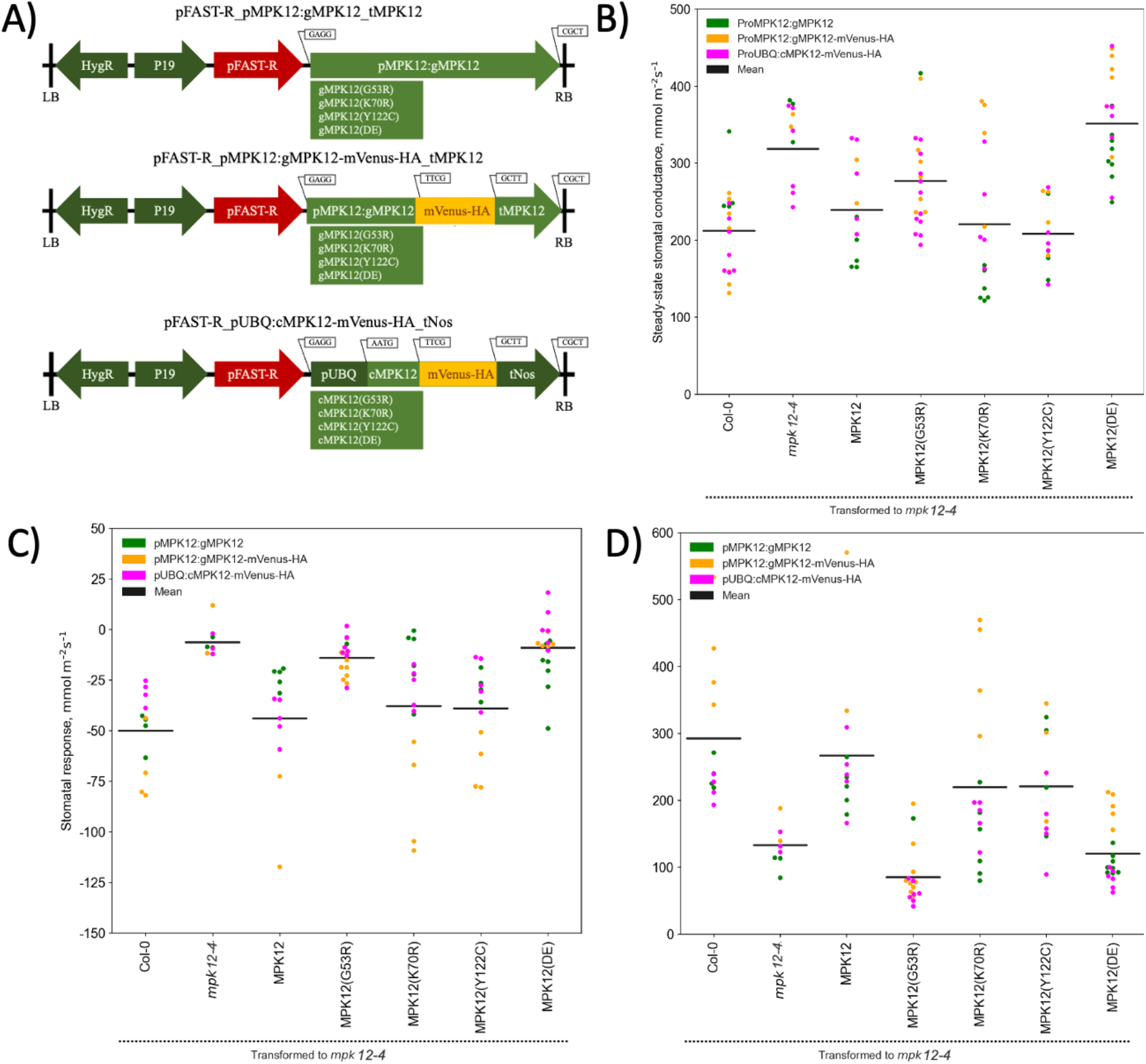
MPK12 variants with K70R and Y122C substitutions, but not the G53R or DE versions, can complement CO_2_ responses in the *mpk12-4* plants. **A)**Diagrams of binary vectors used to transform *mpk12-4* plants: LB- left border T-DNA sequence; RB- right border T-DNA sequence; HygR- Hygromycin B resistance marker gene; P19 – gene silencing suppressor; pFAST-R - *A. thaliana Oleosin* gene fused with a red fluorescence protein sequence; pMPK12 – *MPK12* promoter; gMPK12 – *MPK12* gene sequence with exons and introns; pUBQ – *A. thaliana ubiquitin 10* promoter; cMPK12 – MPK12 coding sequence without introns; mVenus – a yellow fluorescent protein gene; HA - Human Influenza Hemagglutinin tag; tNos - *A. tumefaciens Nopaline synthase* terminator. The overhangs after *Bsa*I digestion of the Golden Gate parts and the PCR products are depicted above the respective parts. **B)** to **D)**Summary of steady state stomatal conductance in **b**), high CO_2_ (from 420 ppm to 800 ppm)-induced closure in **c**), and low CO_2_ (from 420 ppm to 100 ppm)-induced opening in **d**) in T1 plants. The stomatal response is calculated as a difference between the conductance at ambient CO_2_ (420 ppm) before and after 16 minutes of the treatment; dots of different color denote individual T1 plants with binary constructs of the matching color, whereas the black horizontal lines are the average conductance.

**Figure S5.**
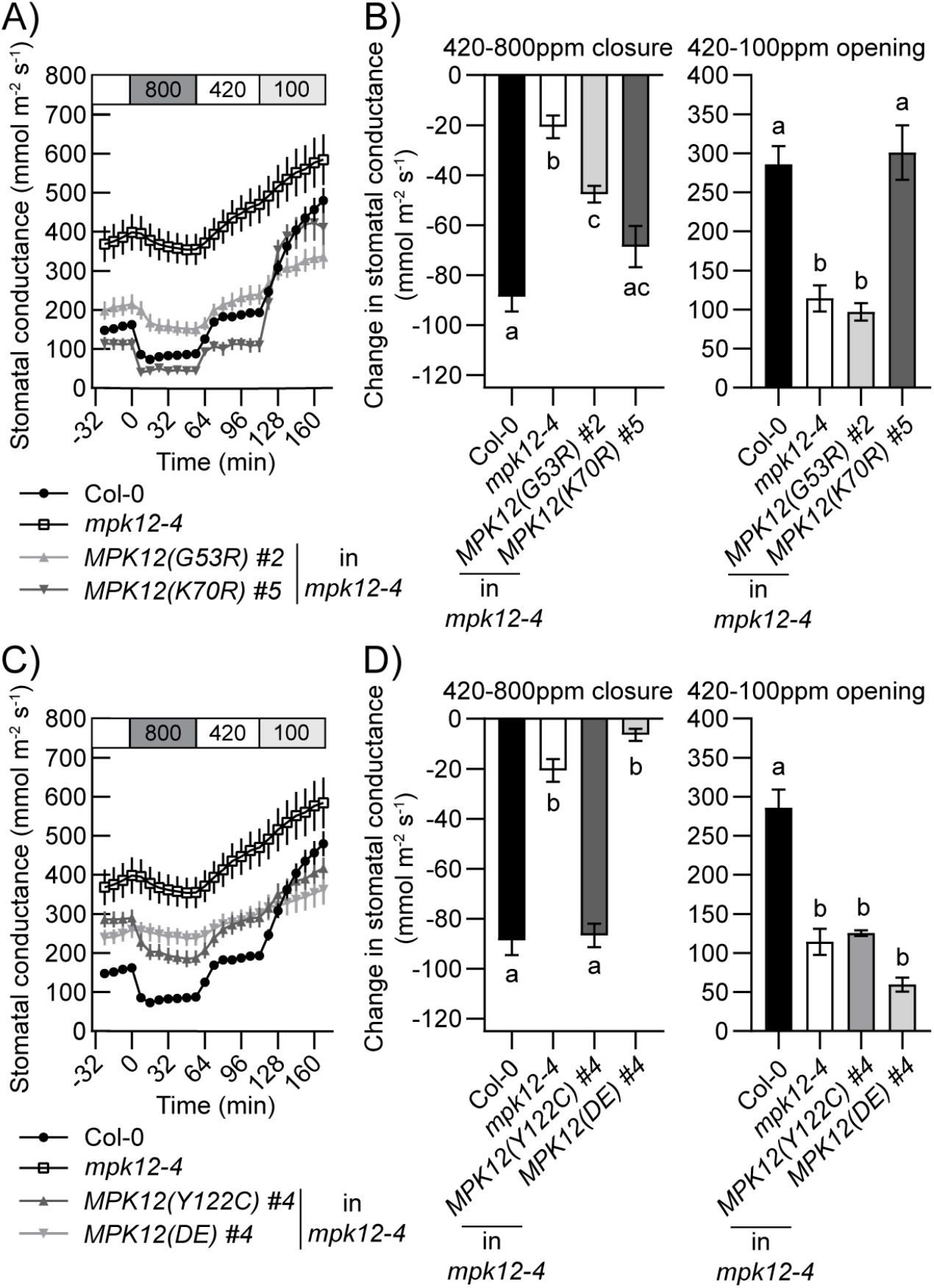
Kinase activity of MPK12 is not required for CO_2_ induced stomatal movements. (This data displays same experiments shown in Fig. 3, but using different T2 lines) **A, C)** Time-resolved patterns of whole-plant stomatal conductance in response to changes in CO_2_ concentrations (in parts per million; ppm) as indicated on top of the panels; intact plants of Col-0 (n=13), *MPK12* deletion line *mpk12-4* (n=12), and *mpk12-4* transformed with *MPK12(G53R)* (n=6), *MPK12(K70R)* (n=5) in **a)**and *MPK12(Y122C)* (n=4), *MPK12(DE)* (n=4) in **c)**were analyzed; data of Col-0 and *mpk12-4* controls are the same as in **Fig. 3a** because they were obtained in the same experimental set. **B, D)**Stomatal closure at 16 min after application of 800 ppm CO_2_ (left panels) and stomatal opening after application of 100 ppm CO_2_ for 56 min (right panels); statistically significant groups are denoted with different letters (ANOVA with Tukey HSD unequal N post hoc test, P < 0.05).

**Figure S6.**
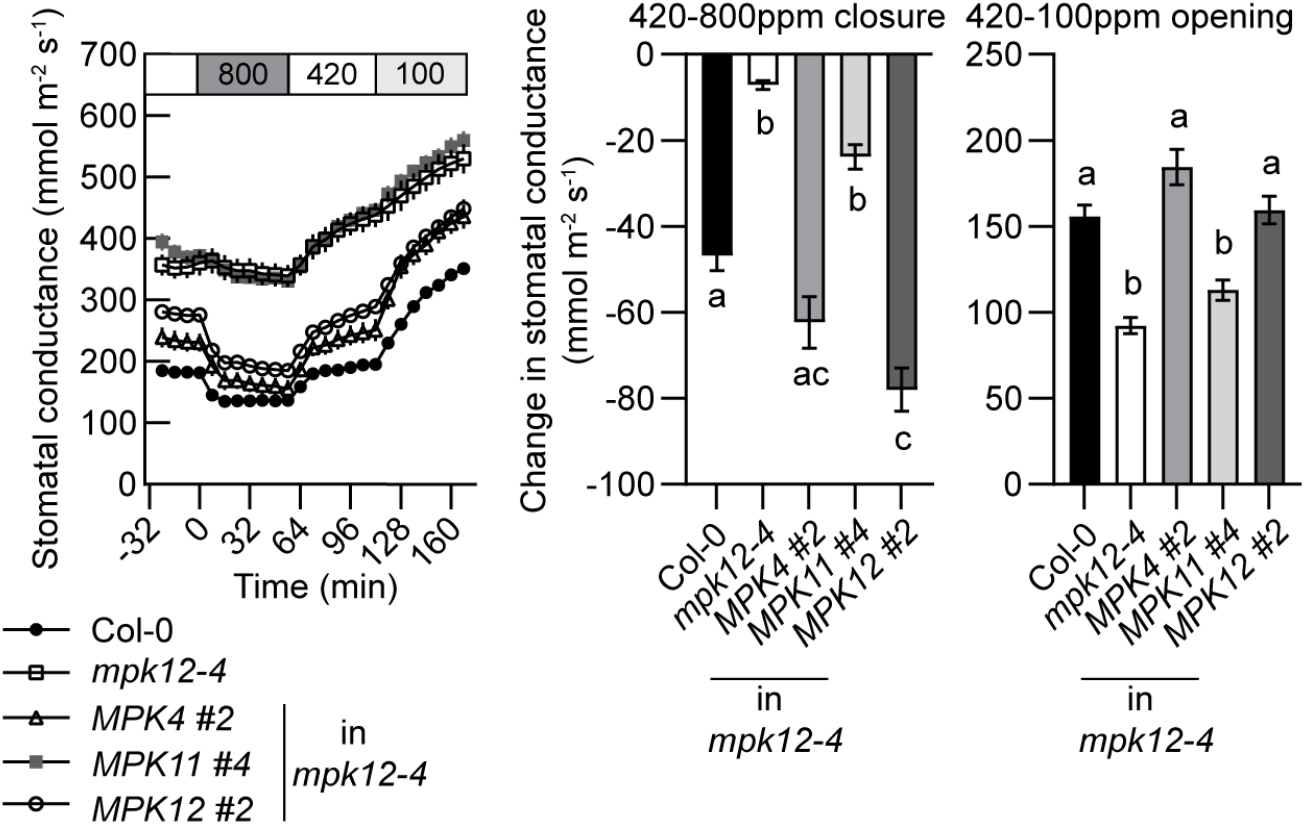
Expression of MPK4 in guard cells can restore mpk12-4 CO_2_ responses but MPK11 cannot. (This data displays the same experiments shown in Fig. 5, but using different T2 lines) **A)** Time-resolved patterns of stomatal conductance in response to changes in CO_2_ concentrations (in parts per million; ppm) as indicated on top of the panels of Col-0 (n=4), *mpk12-4* (n=4) and *mpk12-4* plants complemented with *pMPK12*:*MPK4* (n=5)/*MPK11* (n=6)/*MPK12* (n=5)*-mVenus-HA* constructs; data of Col-0 and *mpk12-4* controls are the same as in **Fig. 5a** because they were obtained in the same experimental set. **B)** Stomatal closure at 16 min after application of 800 ppm CO_2_ (left panel) and stomatal opening after application of 100 ppm CO_2_ for 56 min (right panel); statistically significant groups are denoted with different letters (ANOVA with Tukey HSD unequal N post hoc test, P < 0.05).

## References

Berriri, S., Garcia, A. V., dit Frey, N. F., Rozhon, W., Pateyron, S., Leonhardt, N., Montillet, J.-L., Leung, J., Hirt, H., & Colcombet, J. (2012). Constitutively Active Mitogen-Activated Protein Kinase Versions Reveal Functions of Arabidopsis MPK4 in Pathogen Defense Signaling. The Plant Cell, 24(10), 4281–4293. https://doi.org/10.1105/TPC.112.101253

Clough, S. J., & Bent, A. F. (1998). Floral dip: a simplified method for Agrobacterium-mediated transformation of Arabidopsis thaliana. The Plant Journal, 16(6), 735–743. https://doi.org/10.1046/J.1365-313X.1998.00343.X

Emirick, M. A., Lee, T., Starkey, P. J., Mumby, M. C., Resing, K. A., & Ahn, N. G. (2006). The gatekeeper residue controls autoactivation of ERK2 via a pathway of intramolecular connectivity. Proceedings of the National Academy of Sciences of the United States of America, 103(48), 18101–18106. https://doi.org/10.1073/PNAS.0608849103

Foyer, C. H., & Noctor, G. (2020). Redox homeostasis and signaling in a higher-CO_2_ world. Annual Review of Plant Biology, 71, 157–182. https://doi.org/10.1146/ANNUREV-ARPLANT-050718-095955

Gookin, T. E., & Assmann, S. M. (2014). Significant reduction of BiFC non-specific assembly facilitates *in planta* assessment of heterotrimeric G-protein interactors. Plant Journal, 80(3), 553–567. https://doi.org/10.1111/tpj.12639

Hall, R. A., De Sordi, L., MacCallum, D. M., Topal, H., Eaton, R., Bloor, J. W., Robinson, G. K., Levin, L. R., Buck, J., Wang, Y., Gow, N. A. R., Steegborn, C., & Mühlschlegel, F. A. (2010). CO_2_ acts as a signalling molecule in populations of the fungal pathogen *Candida albicans*. PLoS Pathogens, 6(11). https://doi.org/10.1371/JOURNAL.PPAT.1001193

Hamel, L.-P. P., Nicole, M.-C. C., Sritubtim, S., Morency, M.-J. J., Ellis, M., Ehlting, J., Beaudoin, N., Barbazuk, B., Klessig, D., Lee, J., Martin, G., Mundy, J., Ohashi, Y., Scheel, D., Sheen, J., Xing, T., Zhang, S., Seguin, A., & Ellis, B. E. (2006). Ancient signals: comparative genomics of plant MAPK and MAPKK gene families. Trends in Plant Science, 11(4), 192–198. https://doi.org/10.1016/j.tplants.2006.02.007

Hashimoto-Sugimoto, M., Negi, J., Monda, K., Higaki, T., Isogai, Y., Nakano, T., Hasezawa, S., & Iba, K. (2016). Dominant and recessive mutations in the Raf-like kinase HT1 gene completely disrupt stomatal responses to CO_2_ in *Arabidopsis*. Journal of Experimental Botany, 67(11), erw134. https://doi.org/10.1093/jxb/erw134

Hashimoto, M., Negi, J., Young, J., Israelsson, M., Schroeder, J. I., & Iba, K. (2006). *Arabidopsis* HT1 kinase controls stomatal movements in response to CO_2_. Nature Cell Biology, 8(4), 391–397. https://doi.org/10.1038/ncb1387

Hiyama, A., Takemiya, A., Munemasa, S., Okuma, E., Sugiyama, N., Tada, Y., Murata, Y., & Shimazaki, K. (2017). Blue light and CO_2_ signals converge to regulate light-induced stomatal opening. Nature Communications, 8(1), 1284. https://doi.org/10.1038/s41467-017-01237-5

Hõrak, H., Sierla, M., Tõldsepp, K., Wang, C., Wang, Y.-S., Nuhkat, M., Valk, E., Pechter, P., Merilo, E., Salojärvi, J., Overmyer, K., Loog, M., Brosché, M., Schroeder, J. I., Kangasjärvi, J., & Kollist, H. (2016). A dominant mutation in the HT1 kinase uncovers roles of MAP kinases and GHR1 in CO_2_-induced stomatal closure. The Plant Cell, 28(10), 2493–2509. https://doi.org/10.1105/tpc.16.00131

Hu, S., Xie, Z., Onishi, A., Yu, X., Jiang, L., Lin, J., Rho, H. sool, Woodard, C., Wang, H., Jeong, J. S., Long, S., He, X., Wade, H., Blackshaw, S., Qian, J., & Zhu, H. (2009). Profiling the human protein-DNA interactome reveals ERK2 as a transcriptional repressor of interferon signaling. Cell, 139(3), 610–622. https://doi.org/10.1016/j.cell.2009.08.037

Ichimura, K., Shinozaki, K., Tena, G., Sheen, J., Henry, Y., Champion, A., Kreis, M., Zhang, S., Hirt, H., Wilson, C., Heberle-Bors, E., Ellis, B. E., Morris, P. C., Innes, R. W., Ecker, J. R., Scheel, D., Klessig, D. F., Machida, Y., Mundy, J., … Walker, J. C. (2002). Mitogen-activated protein kinase cascades in plants: a new nomenclature. Trends in Plant Science, 7(7), 301–308. https://doi.org/10.1016/S1360-1385(02)02302-6

Jakobson, L., Vaahtera, L., Tõldsepp, K., Nuhkat, M., Wang, C., Wang, Y.-S., Hõrak, H., Valk, E., Pechter, P., Sindarovska, Y., Tang, J., Xiao, C., Xu, Y., Gerst Talas, U., García-Sosa, A. T., Kangasjärvi, S., Maran, U., Remm, M., Roelfsema, M. R. G., … Brosché, M. (2016). Natural variation in *Arabidopsis* Cvi-0 accession reveals an important role of MPK12 in guard cell CO_2_ signaling. PLOS Biology, 14(12), e2000322. https://doi.org/10.1371/journal.pbio.2000322

Jammes, F., Song, C., Shin, D., Munemasa, S., Takeda, K., Gu, D., Cho, D., Lee, S., Giordo, R., Sritubtim, S., Leonhardt, N., Ellis, B. E., Murata, Y., & Kwak, J. M. (2009). MAP kinases MPK9 and MPK12 are preferentially expressed in guard cells and positively regulate ROS-mediated ABA signaling. Proceedings of the National Academy of Sciences of the United States of America, 106(48), 20520–20525. https://doi.org/10.1073/PNAS.0907205106/SUPPL_FILE/0907205106SI.PDF

Jumper, J., Evans, R., Pritzel, A., Green, T., Figurnov, M., Ronneberger, O., Tunyasuvunakool, K., Bates, R., Žídek, A., Potapenko, A., Bridgland, A., Meyer, C., Kohl, S. A. A., Ballard, A. J., Cowie, A., Romera-Paredes, B., Nikolov, S., Jain, R., Adler, J., … Hassabis, D. (2021). Highly accurate protein structure prediction with AlphaFold. Nature 2021 596:7873, 596(7873), 583–589. https://doi.org/10.1038/s41586-021-03819-2

Kollist, T., Moldau, H., Rasulov, B., Oja, V., Rämma, H., Hüve, K., Jaspers, P., Kangasjärvi, J., & Kollist, H. (2007). A novel device detects a rapid ozone-induced transient stomatal closure in intact *Arabidopsis* and its absence in *abi2* mutant. Physiologia Plantarum, 129(4), 796–803. https://doi.org/10.1111/J.1399-3054.2006.00851.X

Kung, J. E., & Jura, N. (2016). Structural basis for the non-catalytic functions of protein kinases. Structure (London, England: 1993), 24(1), 7. https://doi.org/10.1016/J.STR.2015.10.020

Kushnirov, V. V. (2000). Rapid and reliable protein extraction from yeast. Yeast. https://onlinelibrary.wiley.com/doi/epdf/10.1002/1097-0061%2820000630%2916%3A9%3C857%3A%3AAID-YEA561%3E3.0.CO%3B2-B

Lee, J. S., Wang, S., Sritubtim, S., Chen, J. G., & Ellis, B. E. (2009). *Arabidopsis* mitogen-activated protein kinase MPK12 interacts with the MAPK phosphatase IBR5 and regulates auxin signaling. The Plant Journal, 57(6), 975–985. https://doi.org/10.1111/J.1365-313X.2008.03741.X

Mace, P. D., & Murphy, J. M. (2021). There’s more to death than life: Noncatalytic functions in kinase and pseudokinase signaling. The Journal of Biological Chemistry, 296. https://doi.org/10.1016/J.JBC.2021.100705

Madhani, H. D., Styles, C. A., & Fink, G. R. (1997). MAP kinases with distinct inhibitory functions impart signaling specificity during yeast differentiation. Cell, 91(5), 673–684. https://doi.org/10.1016/S0092-8674(00)80454-7

Marais, D. L. D., Auchincloss, L. C., Sukamtoh, E., McKay, J. K., Logan, T., Richards, J. H., & Juenger, T. E. (2014). Variation in MPK12 affects water use efficiency in *Arabidopsis* and reveals a pleiotropic link between guard cell size and ABA response. Proceedings of the National Academy of Sciences of the United States of America, 111(7), 2836–2841. https://doi.org/10.1073/PNAS.1321429111

Marten, H., Hyun, T., Gomi, K., Seo, S., Hedrich, R., & Roelfsema, M. R. G. (2008). Silencing of NtMPK4 impairs CO_2_-induced stomatal closure, activation of anion channels and cytosolic Ca^2+^ signals in *Nicotiana tabacum* guard cells. The Plant Journal, 55, 698–708. https://doi.org/10.1111/j.1365-313X.2008.03542.x

Matallanas, D., Birtwistle, M., Romano, D., Zebisch, A., Rauch, J., Kriegsheim, A. Von, Kolch, W., von Kriegsheim, A., Kolch, W., Kriegsheim, A. Von, Kolch, W., von Kriegsheim, A., & Kolch, W. (2011). Raf family kinases: Old dogs have learned new tricks. Genes & Cancer, 2(3), 232–260. https://doi.org/10.1177/1947601911407323

Mirdita, M., Schütze, K., Moriwaki, Y., Heo, L., Ovchinnikov, S., & Steinegger, M. (2022). ColabFold: making protein folding accessible to all. Nature Methods 2022 19:6, 19(6), 679–682. https://doi.org/10.1038/s41592-022-01488-1

Mirdita, M., Steinegger, M., Breitwieser, F., Sö Ding, J., & Levy Karin, E. (2021). Fast and sensitive taxonomic assignment to metagenomic contigs. Bioinformatics, 37(18), 3029–3031. https://doi.org/10.1093/BIOINFORMATICS/BTAB184

Patron, N. J., Orzaez, D., Marillonnet, S., Warzecha, H., Matthewman, C., Youles, M., Raitskin, O., Leveau, A., Farré, G., Rogers, C., Smith, A., Hibberd, J., Webb, A. A. R., Locke, J., Schornack, S., Ajioka, J., Baulcombe, D. C., Zipfel, C., Kamoun, S., … Haseloff, J. (2015). Standards for plant synthetic biology: A common syntax for exchange of DNA parts. New Phytologist, 208(1), 13–19. https://doi.org/10.1111/nph.13532

Petersen, M., Brodersen, P., Naested, H., Andreasson, E., Lindhart, U., Johansen, B., Nielsen, H. B., Lacy, M., Austin, M. J., Parker, J. E., Sharma, S. B., Klessig, D. F., Martienssen, R., Mattsson, O., Jensen, A. B., & Mundy, J. (2000). *Arabidopsis* map kinase 4 negatively regulates systemic acquired resistance. Cell, 103(7), 1111–1120. https://doi.org/10.1016/S0092-8674(00)00213-0

Phelan, D. E., Mota, C., Lai, C., Kierans, S. J., & Cummins, E. P. (2021). Carbon dioxide-dependent signal transduction in mammalian systems. Interface Focus, 11(2). https://doi.org/10.1098/RSFS.2020.0033

Shaw, A. S., Kornev, A. P., Hu, J., Ahuja, L. G., & Taylor, S. S. (2014). Kinases and pseudokinases: lessons from RAF. Molecular and Cellular Biology, 34(9), 1538–1546. https://doi.org/10.1128/MCB.00057-14

Shimada, T. L., Shimada, T., & Hara-Nishimura, I. (2010). A rapid and non-destructive screenable marker, FAST, for identifying transformed seeds of *Arabidopsis thaliana*. Plant Journal, 61(3), 519–528. https://doi.org/10.1111/j.1365-313X.2009.04060.x

Sierla, M., Hõrak, H., Overmyer, K., Waszczak, C., Yarmolinsky, D., Maierhofer, T., Vainonen, J. P., Salojärvi, J., Denessiouk, K., Laanemets, K., Tõldsepp, K., Vahisalu, T., Gauthier, A., Puukko, T., Paulin, L., Auvinen, P., Geiger, D., Hedrich, R., Kollist, H., & Kangasjärvi, J. (2018). The receptor-like pseudokinase GHR1 is required for stomatal closure. Plant Cell, 30(11), 2813–2837. https://doi.org/10.1105/tpc.18.00441

Takahashi, Y., Bosmans, K. C., Hsu, P.-K., Paul, K., Seitz, C., Yeh, C.-Y., Wang, Y.-S., Yarmolinsky, D., Sierla, M., Vahisalu, T., McCammon, J. A., Kangasjärvi, J., Zhang, L., Kollist, H., Trac, T., & Schroeder, J. I. (2022). Stomatal CO_2_/bicarbonate sensor consists of two interacting protein kinases, Raf-like HT1 and non-kinase-activity requiring MPK12/MPK4. Science Advances, 8(49). https://doi.org/10.1126/SCIADV.ABQ6161

Taylor, S. S., & Kornev, A. P. (2011). Protein kinases: evolution of dynamic regulatory proteins. Trends in Biochemical Sciences, 36(2), 65–77. https://doi.org/10.1016/J.TIBS.2010.09.006

Tõldsepp, K., Zhang, J., Takahashi, Y., Sindarovska, Y., Hõrak, H., Ceciliato, P. H. O., Koolmeister, K., Wang, Y.-S., Vaahtera, L., Jakobson, L., Yeh, C.-Y., Park, J., Brosche, M., Kollist, H., & Schroeder, J. I. (2018). Mitogen-activated protein kinases MPK4 and MPK12 are key components mediating CO_2_-induced stomatal movements. The Plant Journal, 96(5), 1018–1035. https://doi.org/10.1111/tpj.14087

Tsugama, D., Liu, S., & Takano, T. (2011). A rapid chemical method for lysing *Arabidopsis* cells for protein analysis. Plant Methods, 7(1). https://doi.org/10.1186/1746-4811-7-22

